# Publication practices during the COVID-19 pandemic: Biomedical preprints and peer-reviewed literature

**DOI:** 10.1101/2021.01.21.427563

**Authors:** Yulia V. Sevryugina, Andrew J. Dicks

## Abstract

The coronavirus pandemic introduced many changes to our society, and deeply affected the established in biomedical sciences publication practices. In this article, we present a comprehensive study of the changes in scholarly publication landscape for biomedical sciences during the COVID-19 pandemic, with special emphasis on preprints posted on bioRxiv and medRxiv servers. We observe the emergence of a new category of preprint authors working in the fields of *immunology, microbiology*, *infectious diseases*, and *epidemiology*, who extensively used preprint platforms during the pandemic for sharing their immediate findings. The majority of these findings were works-in-progress unfitting for a prompt acceptance by refereed journals. The COVID-19 preprints that became peer-reviewed journal articles were often submitted to journals concurrently with the posting on a preprint server, and the entire publication cycle, from preprint to the online journal article, took on average 63 days. This included an expedited peer-review process of 43 days and journal’s production stage of 15 days, however there was a wide variation in publication delays between journals. Only one third of COVID-19 preprints posted during the first nine months of the pandemic appeared as peer-reviewed journal articles. These journal articles display high Altmetric Attention Scores further emphasizing a significance of COVID-19 research during 2020. This article will be relevant to editors, publishers, open science enthusiasts, and anyone interested in changes that the 2020 crisis transpired to publication practices and a culture of preprints in life sciences.

## Introduction

The lifecycle for any research starts and ends with a scholarly communication. Despite a variety of avenues to communicate research findings, the foundation of the modern publication practices is a publication in a peer-reviewed journal. The peer-review system is, at present, deeply engraved in scientific minds as the golden standard for research quality. Certainly, the peer-review process improves the drafted manuscript, but previous studies showed that its positive effect on the overall quality of the final report is minor [1]. Besides, the traditional peer-review system is notorious for reviewer bias, lack of agreement between reviewers, harsh criticism concealed by anonymity, multiple cycles of reviews and rejections by different journals, and associated delays and expenses [2].

Alternatively, or additionally, authors may choose to deposit their manuscripts to preprint servers, institutionally or privately supported repositories of preprints. A preprint is a complete manuscript shared publicly prior to officially undergoing the peer-review process. Notably, it has likely undergone many rounds of internal review by authors and colleagues involved in its preparation to ensure that ideas presented are well-versed and that the study can be easily validated by the supporting data. A preprint submitted to one of the main preprint servers undergoes basic screening for its scope, plagiarism, ethical issues and compliance, often performed as quickly as within 24 hours, after which it is published online with a digital object identifier (DOI) that allows it to be citable and trackable. Once posted on a preprint server, preprint can be read, commented on-site or by email, and further shared on the Web and through social media (e.g. Twitter). Additionally, any revision of a preprint’s content or status, such as publication in a peer-reviewed journal, is time-stamped.

Over the last decade, biomedical sciences have been slowly adapting a preprint culture. The bioRxiv server owned by Cold Spring Harbor Laboratory was launched in November 2013 with the purpose of covering all aspects of life sciences research [3], and by 2018, it accumulated 37,648 preprints. By the end of September 2020, this number has doubled to 97,194 preprints. The medRxiv preprint server, launched in June 2019 by BMJ, Yale University, and Cold Spring Harbor Laboratory to cover all aspects of research in the medical, clinical, and related health sciences [4], contained 11,329 preprints on September 30, 2020. The first and the most well-known preprint server, arXiv.org was launched in 1991, and by the end of September 2020 had accommodated 1,769,336 preprints [5] in the fields of physics, mathematics, computer science, quantitative biology, quantitative finance, statistics, electrical engineering and systems science, and economics. The preprint platforms were widely explored as hubs for disseminating scientific findings [6], especially for early-stage researchers [7], which revealed a number of social and technical issues associated with their adoption [8]. The top four concerns with preprint servers included scooping of scientific results [9], poor data quality [10], spread of misinformation [11], and varied and non-transparent deposition policies [6]. Additionally, many authors refrained from depositing preprints due to unclear journal policies on whether a preprint will be accepted for a journal publication after being deposited on a server [12]. Nevertheless, research communities, including life sciences [13], persisted in their adoption of preprints [14], gradually realizing that a preprint submission offers many advantages:

1. Allows authors to establish the scientific priority by timestamping the first public record of the research study [15];
2. Provides authors with a community feedback that can help them to further improve the manuscript quality [16, 17];
3. Expedites research sharing [15, 16];
4. Increases research visibility [18, 19];
5. Streamlines the journal submission process [20];
6. Allows sharing studies that are difficult to publish in traditional journals (works-in-progress, negative results, replications, contradictions) [16];
7. Provides an open-access publication record [16].

Those benefits are what make scientific publishers and funders embrace preprints and acknowledge them as works of scholarly communication. In June 2020, the National Institutes of Health (NIH) launched the Preprint Pilot [21] to index preprints that come from NIH-funded projects in PubMed, the largest database of biomedical scholarly works. This pilot builds on the role of PubMed Central as a repository for NIH-supported [22] peer-reviewed articles and NIH’s guide notice NOT-OD-17-050 [23]. Library system vendors such as EBSCO, Proquest, Ex Libris, and OCLC WorldCat have been investing substantially in open access discovery tools for a variety of open sources, including preprints [24]. The practice of inviting journal submissions from preprint servers is wide spread [25]. In June 2020, MIT Press and the Berkeley School of Public Health launched a new COVID-19 journal, *Rapid Reviews: COVID-19*, which, after a thorough peer-review, will publish preprint articles with good research and discredit those with bad [26]. Considering the reported influence of preprints on policy-making during the COVID-19 pandemic [27] and ongoing article retractions [28], the concern about the quality of un-refereed preprints is genuine [29]. The Sinai Immunology Review Project is an example of an institutional effort to review and validate the COVID-19 related preprints posted to medRxiv and bioRxiv servers [30]. The Review Commons platform, launched in December 2019, is another initiative on that direction, where preprint authors are offered an opportunity to request a journal-independent portable peer review [31]. Lastly, it is worth mentioning new editorial policies from *eLife* that resulted in the launch of Preprint Review [32] and made the preprint deposition mandatory prior to a journal submission [33].

Since the first report on a pneumonia of unknown origin in Wuhan, China on Dec 30, 2019 [34], over 10,000 articles have been published in scholarly journals and on preprint servers, as well as about 4,000 clinical trials registered. These figures attest an outstanding response of the scientific community to what was in January 2020 simply known as the “novel coronavirus” [35]. As our health workers fought on the front lines against coronavirus and general population followed rapidly changing policies, the scientific community immediately sought to contribute by sharing and exchanging newly discovered information promptly and openly. Fortunately, the necessary tools were already in-place and biomedical researchers quickly adopted preprint servers as suitable platforms for timely, open, and transparent scientific communication. In this paper, we set our goal to assess all aspects related to this adoption by analyzing the effect of COVID-19 pandemic on publication practices in biomedical sciences.

## Methods

### Scope

The scope of this study is COVID-19 related literature. For preprints, we focused on medRxiv and bioRxiv preprint servers that compiled all their COVID-19 SARS-CoV-2 preprints in the database that at the time of this study contained 1,964 bioRxiv and 7,258 medRxiv preprints. For peer-reviewed scholarly works, we referred to journal and review articles indexed in PubMed, of which about 1,400 had associated preprints.

### Timeline

The first media statement on ‘viral pneumonia’ in Wuhan, People’s Republic of China appeared on Dec 31, 2019 [36]. By the end of January, the first research reports appeared as preprints, clinical trials, and journal articles [37]. In this manuscript, we focus on preprints deposited between Jan 1 and Sept 30, 2020. The publication rates for these preprints were evaluated on Sept 30 and on Dec 7, 2020. The deposition rate for preprints was evaluated during Jan 1 – Nov 30, 2020. Altmetric analysis is based on articles published during Jan 1 - Nov 19, 2020, and includes articles with a publication date of Dec 1, 2020 (retrieved on Nov 19, 2020). The latter date was included because we noticed that when date is unknown, Dimensions automatically assigns it the first of the month. Similar enrichment of dates on the first of the year and each month was observed in Crossref [38].

### Terminology

*Preprint* is defined according to the COPE (Committee of Publication Ethics) definition [39]: “A preprint is a scholarly manuscript posted by the author(s) in an openly accessible platform, usually before or in parallel with the peer review process.” As the main subjects of this paper are COVID-19 related preprints posted on bioRxiv and medRxiv preprint servers, we will often address to them as simply “preprints”. When discussing other preprints, preprint servers, or publication topics, we will specify it in the text.

*Preprint server* – a repository of preprints. This study specifically focuses on bioRxiv and medRxiv preprint servers.

*Preprint category* – a subject-filed category defined by a preprint server and selected by authors during the deposition process.

*Journal category* – a category assigned by Scopus to the journal to define its scope.

*Published preprint* – a preprint of an article in a peer-reviewed journal.

*Elapsed time* (*T*_Σ_) – interval between the date when a preprint was deposited to the server and publication date for its journal article analogue.

*Pre-submission time* (*tα*) – interval between the date when a preprint is deposited to the server and the date when it is submitted to the journal.

*Review time* (*t_R_*) – interval between the date when manuscript is submitted to the journal and the date it is accepted for publication.

*Production stage time* (*tβ*) – interval between the acceptance date for a manuscript and the date the peer-reviewed journal article appears online.

### Data sources

This paper examines data acquired from a number of sources, including the database of COVID-19 SARS-CoV-2 preprints from medRxiv and bioRxiv [40], Rxivist [41], Crossref [42], E-utilities [43], Dimensions [44], CORD-19 [45], and CADRE[46].

Metadata for each individual COVID-19 preprint deposited to bioRxiv or medRxiv was gathered by accessing the bioRxiv database of COVID-19 SARS-CoV-2 preprints from medRxiv and bioRxiv, to which we will further refer as BioRxiv API [40]. Data were retrieved in JavaScript Object Notation (JSON) format. Data analysis and visualization was done in Python (pandas, numpy, requests, matplotlib, bokeh, and seaborn) using Jupyter Notebook.

To search PubMed, we used Entrez Programming Utilities (E-utilities) [43], an application programming interface (API) that allows searching 38 databases from the National Center for Biotechnology Information (NCBI). For E-Utilities, data were downloaded *via* CSV and converted to Microsoft Excel for further analysis and visualization.

Rxivist [41] is a Python-based web crawler that parses the bioRxiv website, detects newly posted preprints, and stores metadata about each item in a PostgreSQL database. The metadata we extracted contained title, authors, submission date, category, DOI for preprint and, if published, the new DOI and the journal of publication.

Crossref [42] is an official DOI registration agency of the International DOI Foundation that establishes a cross-publisher citation linking system for academic that include journals, conference proceedings, books, data sets, etc. It works with thousands of publishers to provide authorized access to their metadata including DOI, publication date and other basic information.

CORD-19 or COVID-19 Open Research Dataset [47] is a free resource of over 200,000 scholarly articles about COVID-19, SARS-CoV-2, and related coronaviruses prepared by the Allen Institute for AI (AI2) in collaboration with many partners and released on March 16, 2020. We used its 2020.09.02 release downloaded on 2020.09.30 from CADRE [46] for metadata associated with refereed journal articles.

Dimensions [48] is a comprehensive database that links scholarly outputs to a research analytics suite to track the impact of research across its life cycle. Dimensions tracks many preprint servers [49] but we only used it for bioRxiv and medRxiv preprints (see Data Flow Chart in SI for the links between used datasets).

### Statistical analysis

Descriptive analysis of the data, Student’s t-test, and a one-way ANOVA were conducted on the Statistical Package for Social Sciences version 27 (SPSS). Data are presented as means (M) ± standard deviations (SDs), accompanied by medians and modes, when necessary.

### Altmetric data

Altmetric Attention Scores were retrieved from Dimensions by querying for articles published between Jan 1 and Nov 19, 2020, in selected journals. For COVID-19 related publications in a journal, we used the recommended query for COVID-19 [50] supplemented with the query for journal id. For journal publications unrelated to COVID-19, we gathered all articles from the journal and removed those related to COVID-19 from the dataset. For COVID-19 articles that were associated with preprints, we matched DOI’s of articles found by Dimensions to DOI’s of articles we identified earlier as published preprints. To identify articles that were not associated with preprints, we removed from the dataset of COVID-19 related publications all articles that had associated bioRxiv and medRxiv preprints.

### Data challenges and study limitations

#### Analysis of published preprints

When a preprint is published in a peer-review journal, a reference to the new DOI of the journal article appears next to its title, and DOIs of a preprint and a published article are permanently linked in indexing platforms and tools, which pull from various APIs. Rxivist [41] showed to be an excellent tool for extracting published DOIs for preprints eventually appearing as peer-reviewed journal articles but only when bioRxiv records linked preprints to their external publications. Rxivist also had a two weeks delay in updating its metrics, and it might be of this delay that some peer-reviewed preprint analogues were missing from Rxivist. Additionally, at the time of our study, Rxivist did not include medRxiv preprints in its database, which changed after Nov 27, 2020. We found that the most reliable method of extracting metadata about each individual preprint was by accessing the BioRxiv API [40]. Using the Python library requests, we were able to extract information about each preprint based on DOI, which gave us a column called ‘published.’ Within this column, if the preprint was also published in a journal, the metadata provided the DOI that corresponded to the published version of the paper. To ensure we found all published preprints, we also accessed data from Crossref, Dimensions, and CORD-19 APIs. To establish the linkage between the preprints and corresponding peer-reviewed journal articles we performed both, DOI and title matching. All channels were then combined and duplicates were dropped. For detailed demonstration of data obtained by every data channel, see Published Collections in SI.

To validate whether we found all peer-reviewed preprint versions based on a combination of Rxivist, Crossref, CORD-19, Dimensions, and BioRxiv API, we randomly selected a sample of 100 preprints that our data returned as “unpublished” from both bioRxiv and medRxiv, and searched Google Scholar by title. Our analysis of “unpublished” preprints returned 10% of bioRxiv and 4% of medRxiv preprints as being published in refereed journals. All found journal publications had slight modifications in article titles or authors’ list, and the original “unpublished” preprints were not linked on preprint servers to the corresponding published versions. In comparison, this false-negative rate is lower than the 37.5%, reported by Blekhman *et al*. [51]. All manually found journal article versions of “unpublished” preprints were manually added to data discussed in this article.

### Double DOI

When we looked for published preprints based on title matching, we encountered a few instances when two published DOIs existed for a peer-reviewed preprint version. In one case, it was erratum for the paper and in the other case it was a publication on another preprint server. In both cases, we used only the DOI for the article in the peer-reviewed journal and publication on another preprint server was removed from further analysis. We also encountered a few cases when preprints with different DOIs were linked to the same DOI of the published version. On inspection, preprints with different DOIs were somewhat similar in titles and authors’ list but not identical. For our analysis, we kept only one DOI for a preprint that was published earlier.

### PubMed

As mentioned in the Introduction, the NIH Preprint Pilot started in June 2020 and at this stage, it primarily focuses on NIH-supported and COVID-19 related preprints from various servers. By Sept 26, PubMed indexed 1,048 preprints from medRxiv, bioRxiv, ChemRxiv, arXiv, Research Square, and SSRN, of which 1,043 were on COVID-19, and this constituted only 11.5% of 9,072 medRxiv and bioRxiv COVID-19 related preprints from the BioRxiv API. For these reasons, we did not use PubMed as a data source for preprints. We used PubMed (through E-Utilities) to obtain metadata on peer-reviewed articles of “Journal Article” and “Review” article types as the most traditional types of scholarly output. These two types constitute about 24% [52] of all PubMed publications that include 187 different publication types [53].

In analyzing PubMed dates, we found that articles with a missing day-of-publication were coded as being published on January 1^st^; a similar issue was reported earlier for Crossref dates [38]. Based on low number of preprints in January, we decided to avoid discussing January data for PubMed (this month is omitted in Fig 2).

### Categories

In general, we used a single category for a preprint as indicated in metadata from the BioRxiv API. However, as of September 25, we found six out of total 1,956 of COVID-19 related bioRxiv preprints (0.3%) that displayed two categories. Since this contradicted the servers’ statement that “Only one subject area can be selected for an article”, we omitted the additional category in our analysis. The journals’ scope categories were extracted from Crossref [54].

### Publication process delays

Preprints deposition dates were extracted from Crossref. For journal articles received, accepted, and published online dates, we used E-Utilities: PubMedPubDate@PubStatus = “received”; PubMedPubDate@PubStatus = “accepted” and ArticleDate@DateType=“Electronic”. When ArticleDate@DateType=“Electronic” from PubMed was not available, we substituted it with the “created-date” from Crossref.

Before deciding on which dates to use in our studies, we carefully analyzed those used in previous studies and noted some inconsistency between different authors (Table 4). Unfortunately, dates from different sources intended to represent the article publication date differ between each other (see Article Publication Dates in SI) and are often incomplete. The amount of missing dates in our dataset was 44.7% for “pub_date” from PubMed and over 90% for “published-online” or “published-print” from Crossref. In this article, we referred to refereed journals to resolve disputable publication dates. Our scrupulous analysis showed that ArticleDate@DateType=“Electronic” from PubMed is the date that exactly corresponds to the online publication date of a journal article. When this date was not available (6.6% of the cases), we used the “date-created” from Crossref that indicates the day when DOI for the article is created. The “date-created” from Crossref differed from the online publication date of a journal article in 24% of the cases. This number was only 15.4% for “pub_date” from PubMed, but the latter had high percentage of missing dates, while the “date-created” from Crossref had none. These variations in measuring article publication dates may lead to some discrepancy between reports and inconsistent statistical analysis. Based on our experience, we recommend extracting article publication dates either directly from the journal articles or using ArticleDate@DateType=“Electronic” in PubMed and/or “date-created” from Crossref.

To assess the preprint *pre-submission time*, we subtracted the preprint deposition date from the date the journal articles was “received”. To assess the *review time*, we subtracted the date the journal articles was “received” from the date it was “accepted”. To assess the *production stage time*, we subtracted the date the journal article was “accepted” from the date it was posted online (“Electronic”). To evaluate the *elapsed time*, we subtracted the preprint deposition date from the date the journal articles was posted online. One COVID-19 related journal article [55] was excluded from our analysis of publication delays due to a peculiar date of submission, in May 2019, long before the emergence of the first report on COVID-19.

## Results

### 1. Submission rates during the coronavirus pandemic

With the COVID-19 outbreak, both bioRxiv and medRxiv experienced a surge of COVID-19 preprint submissions. The first preprint on COVID-19 appeared on Jan 13 of 2020 [56], and the number of submissions increased at a staggering rate afterwards (Fig 1). By Dec 4, 2020, the number of COVID-19 preprints reached 8,750 on medRxiv and 2,416 on bioRxiv. In comparison, we identified only 187 Zika related, and 17 Ebola related preprints deposited to bioRxiv during the entire duration of each epidemic (for Zika: Nov 2015 - Aug 2017; for Ebola: May 2014 - Jan 2016).

**Fig 1.**
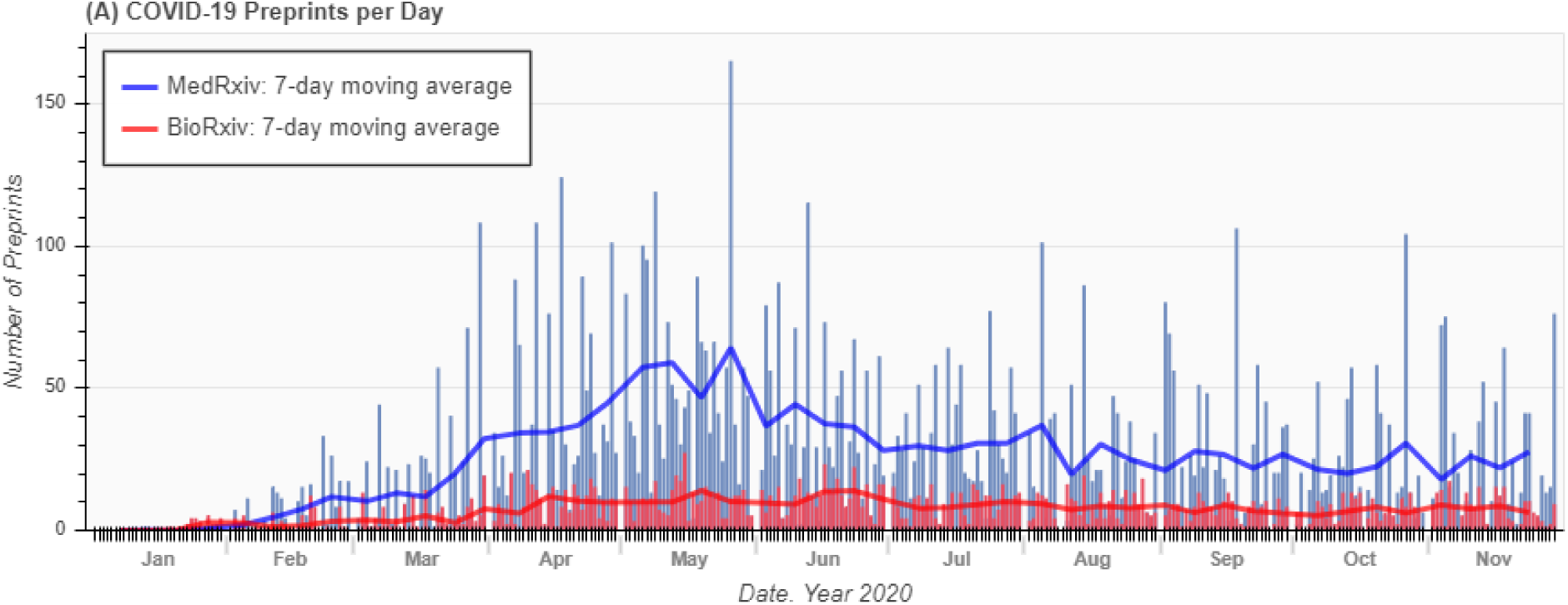
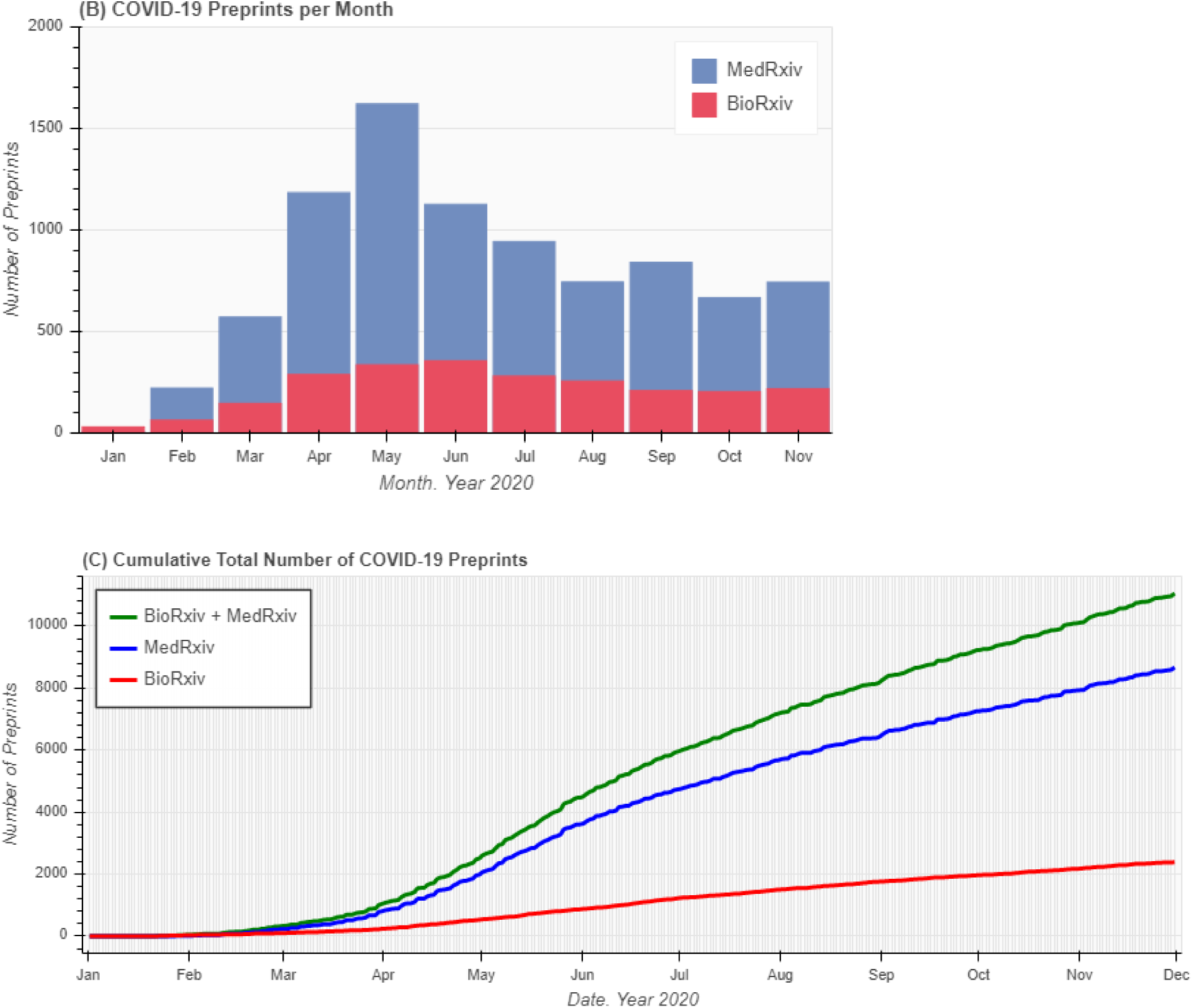
Trends in submission of COVID-19 preprints. COVID-19 Preprints, bioRxiv (red), medRxiv (blue), and combined (green): (A) Daily; (B) monthly; (C) cumulative during Jan 1 – Nov 30, 2020.

Consistently throughout the pandemic, medRxiv experienced a significantly higher flux of COVID-19 preprints as compared to bioRxiv (Table 1 and Scholarly Output in SI). On average, medRxiv preprints on COVID-19 constituted 78% (SD = 2%) of combined bioRxiv and medRxiv preprints on any single month, except January, when the number of COVID-19 related medRxiv preprints was only 27% of COVID-19 related bioRxiv preprints. May was the most productive month for authors of medRxiv preprints. In June, the number of medRxiv COVID-19 preprints declined by 31%, while the number of bioRxiv preprints increased by 6%. After June, we noted a slow decline in the number of COVID-19 preprints on both servers, with the total number of bioRxiv and medRxiv COVID-19 preprints in November representing only 49% of those deposited in May.

**Table 1.**
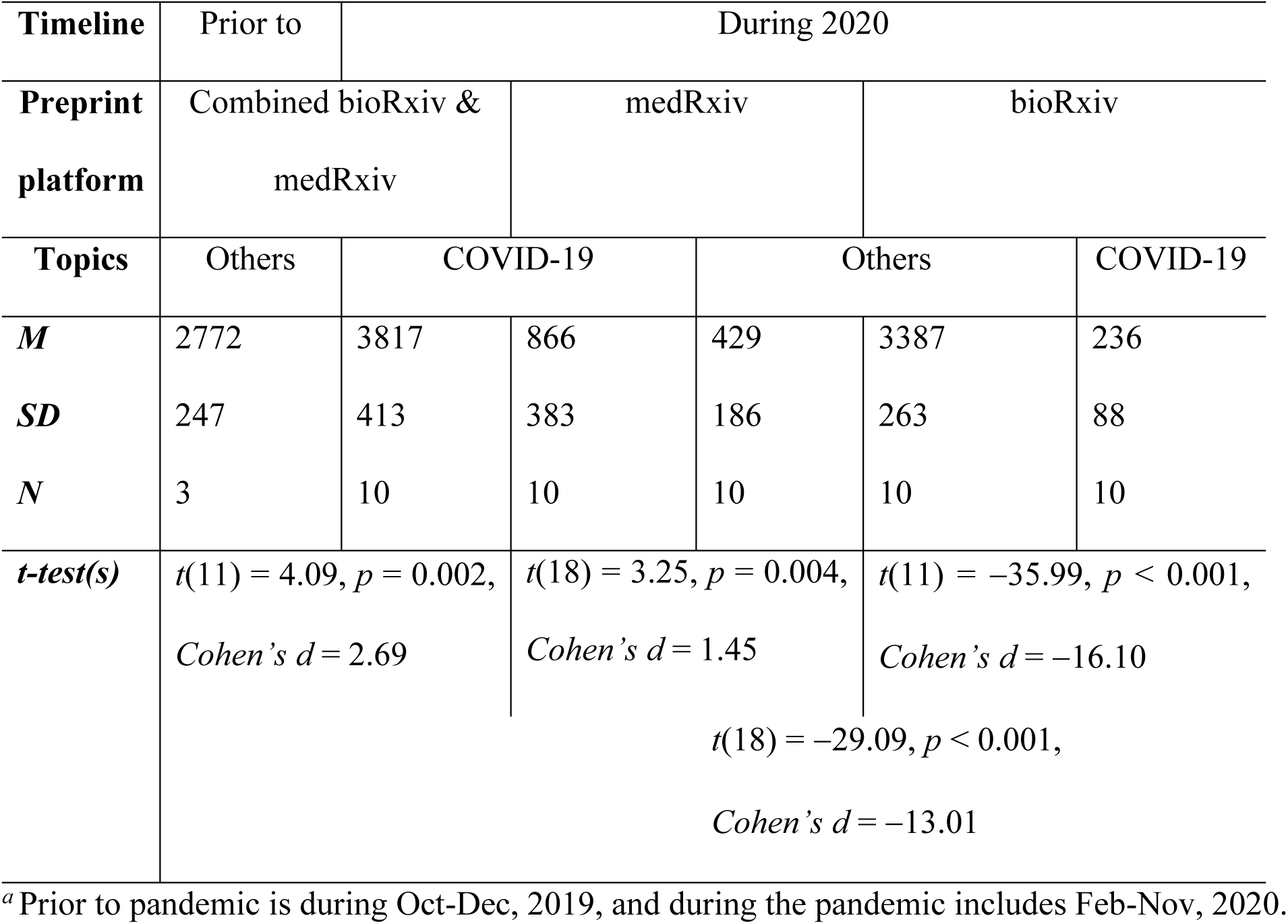
Descriptive statistics and independent samples t-test(s) for preprint submissions prior to and during the COVID-19 pandemic.*^a^*

We then analyzed how the high flux of COVID-19 preprints compares to that of non-COVID-19 preprints and biomedical journal articles published during the same period (Fig 2, Table 1). Notably, during the pandemic, we observe the growth in preprints submissions not only on COVID-19 but also on other topics unrelated to coronavirus. Consistently, bioRxiv received significantly more of non-COVID-19 preprints than medRxiv. While the COVID-19 submissions have been slowly declining since June, those unrelated to coronavirus are relatively stable for bioRxiv and have been increasing for medRxiv (see Scholarly Output in SI), indicating a growing popularity of medRxiv server.

**Fig 2.**
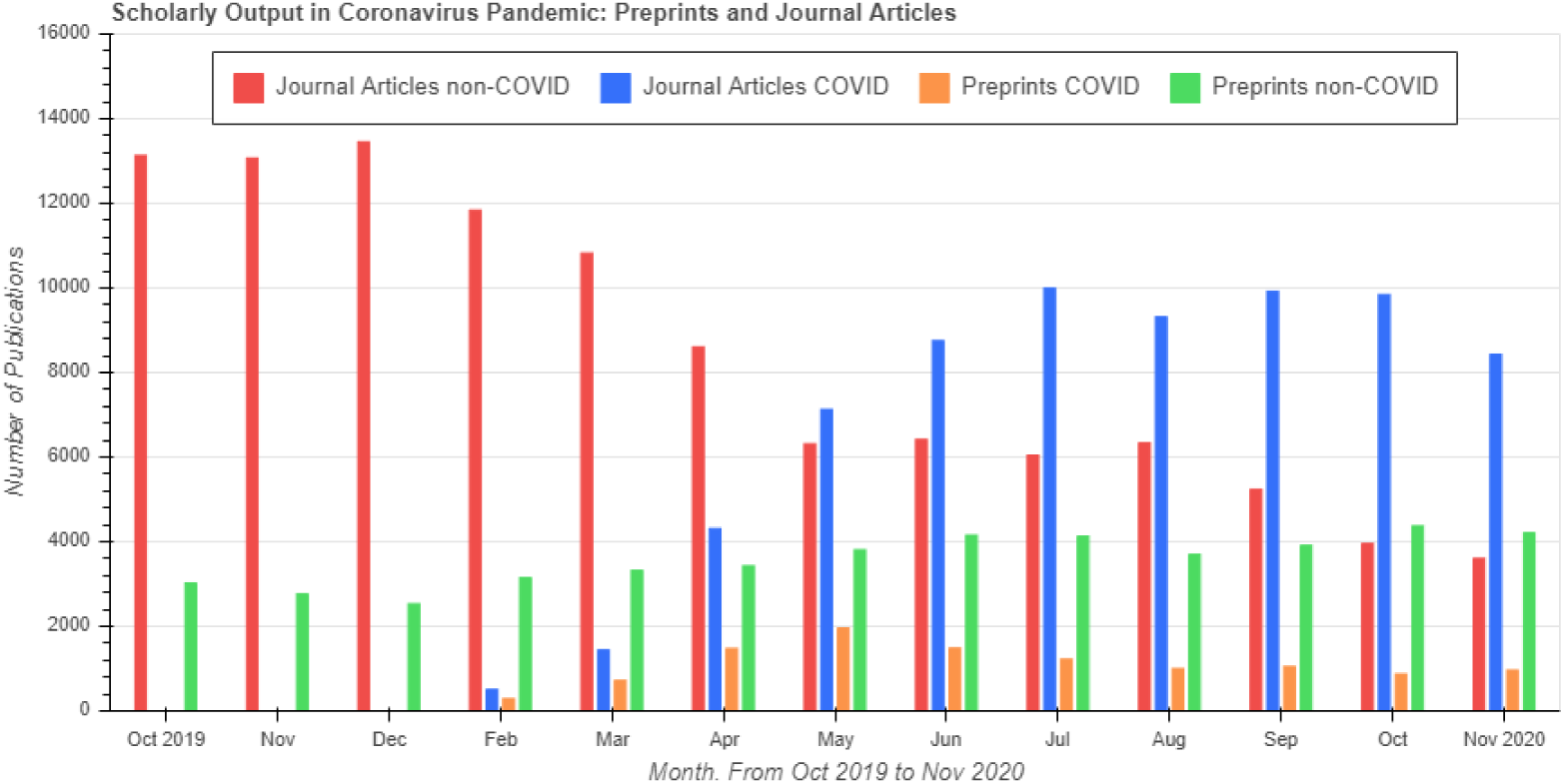
Scholarly output during the coronavirus pandemic. Combined medRxiv and bioRxiv preprints related to COVID-19 (orange); combined medRxiv and bioRxiv preprints unrelated to COVID-19 (green); PubMed journal articles related to COVID-19 (blue); PubMed journal articles unrelated to COVID-19 (red), prior to (Oct – Dec, 2019) and during (Feb – Nov, 2020) the coronavirus pandemic.

The growth of preprints, however outstanding, is lagging behind the amount of peer-reviewed journal articles (see Scholarly Output in SI). In 2019, preprints represented 2.6% of all biomedical literature in PubMed [57]. This percentage was derived with respect to biomedical preprints from multiple servers and all biomedical literature indexed in PubMed [58], while our study is defined by bioRxiv and medRxiv preprints in relation to “Journal Article” and “Review” article types in PubMed. Based on our analysis, in February, the amount of COVID-19 preprints from medRxiv and bioRxiv constituted only 2% of biomedical articles on all topics but this fraction increased to 15% in May (Fig 3). The number of peer-reviewed articles on COVID-19 has been growing since the start of pandemic reaching a peak in July. In contrary, the number of unrelated to coronavirus peer-reviewed literature has been slowly declining. As a result, the fraction of COVID-19 journal articles with respect to all articles indexed in PubMed has been increasing since the start of pandemic and reached 71% in October. At that time, the amount of COVID-19 bioRxiv and medRxiv preprints was at 9% with respect to COVID-19 peer-reviewed literature in PubMed, but this fraction was as high as 57% in February 2020. Thus, early in pandemic, there were over half as many preprints as there were peer-reviewed articles about the newly emerged coronavirus.

**Fig 3.**
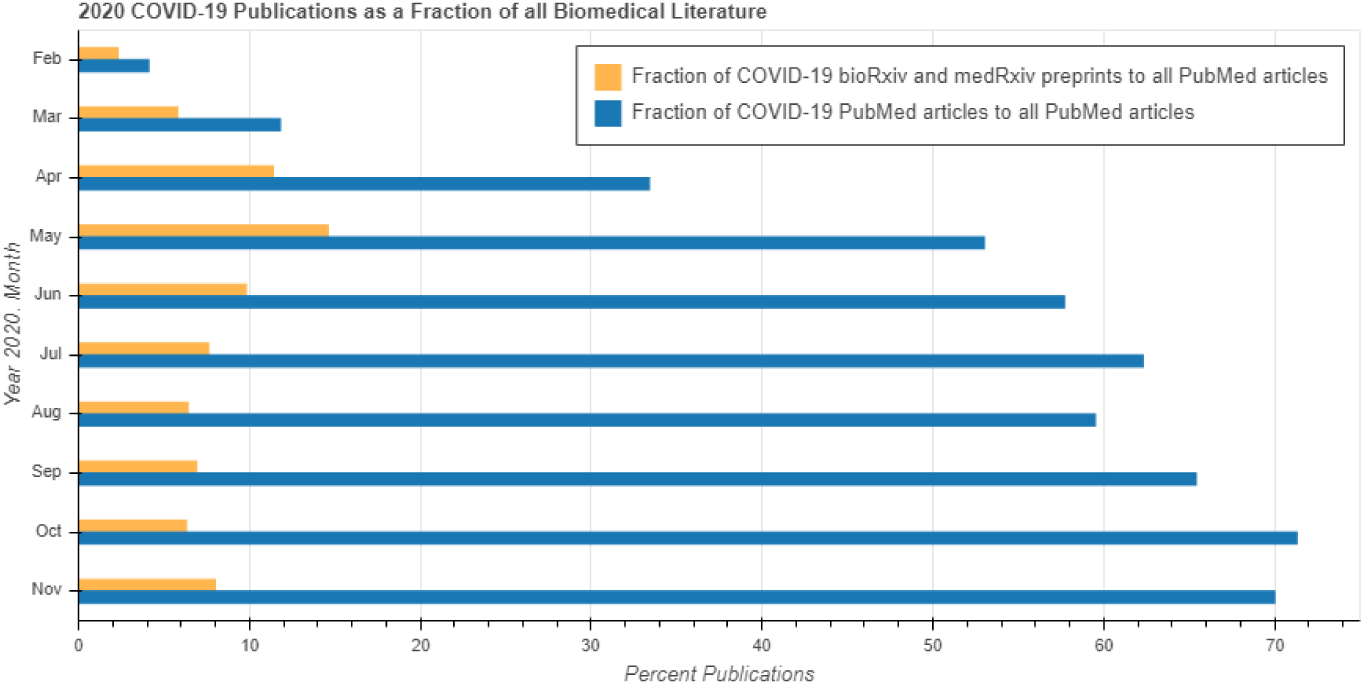
Percent ratio of COVID-19 publications to all journal articles in PubMed. Combined COVID-19 medRxiv and bioRxiv preprints (orange) and COVID-19 journal and review articles (blue).

### 2. Preprints categories

Preprints are deposited in a certain category that defines their subject area. As expected, *microbiology*, the field that studies microscopical organisms, such as viruses, is the top category for bioRxiv preprints on COVID-19 (Fig 4A). Other leading bioRxiv categories for COVID-19 preprints are *bioinformatics* that analyzes and interprets large and complex biological data, and *immunology* that studies the immune system. The top three leading categories for COVID-19 medRxiv preprints are *infectious diseases*, *epidemiology*, and *public and global health* (Fig 4B). Clearly, the study of *infectious diseases* focuses on infectious agents, e*pidemiology* deals with the possible control of diseases, and *public and global health* focuses on disease prevention, all of them closely related to our understanding of the nature of coronavirus, and the development of corresponding diagnostic methods and treatment. The three leading categories for COVID-19 preprints in bioRxiv incorporate 1.3 times as many preprints as the remaining 20 subject categories. The three leading categories for COVID-19 preprints in medRxiv incorporate 3.2 times as many preprints as the remaining 48 subject categories (see Categories Analysis in SI). As expected, the leading categories have the highest daily accumulation rates (Fig 5).

**Fig 4.**
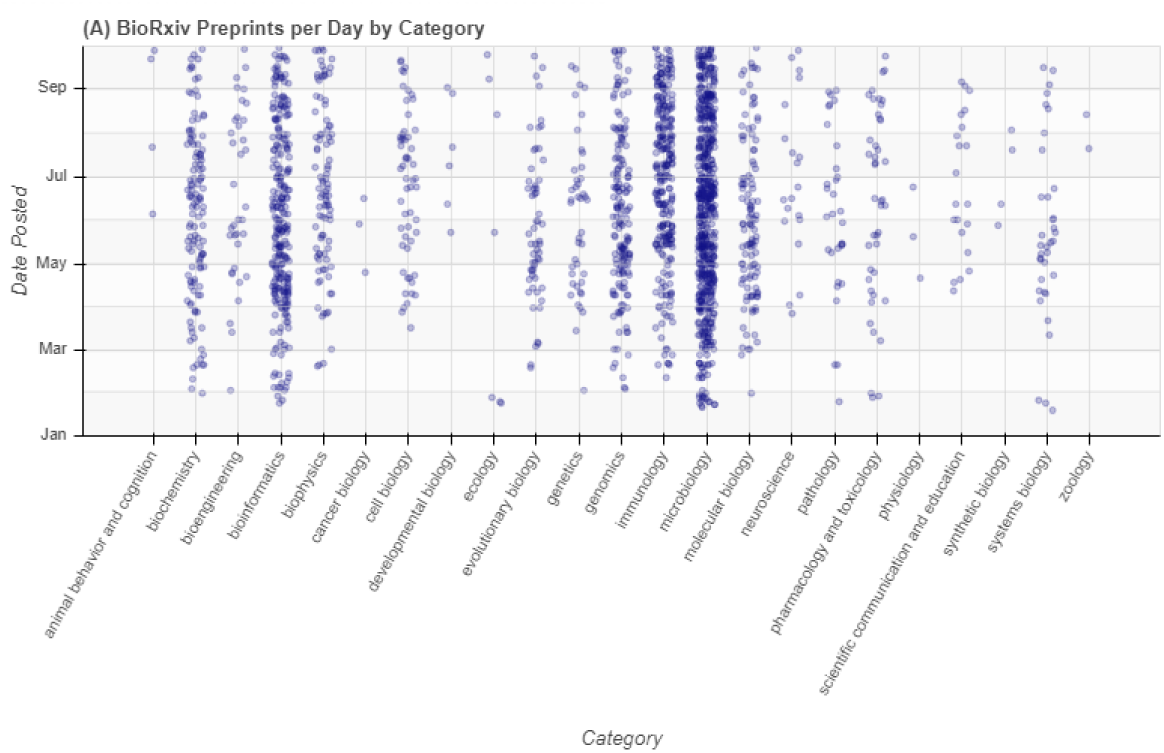
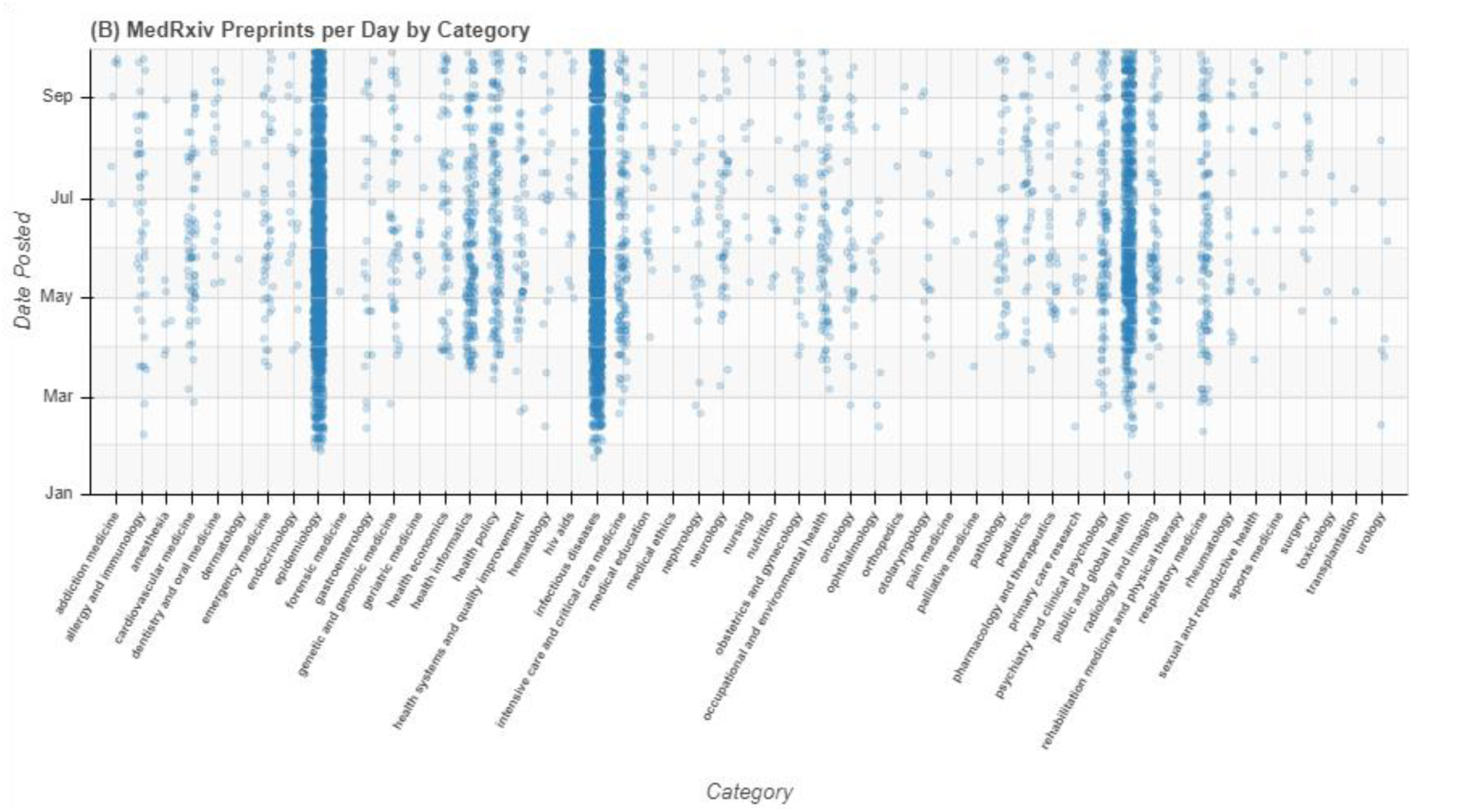
Daily COVID-19 preprint submissions by preprint category. (A) BioRxiv and (B) medRxiv preprint submissions (represented by dots).

**Fig 5.**
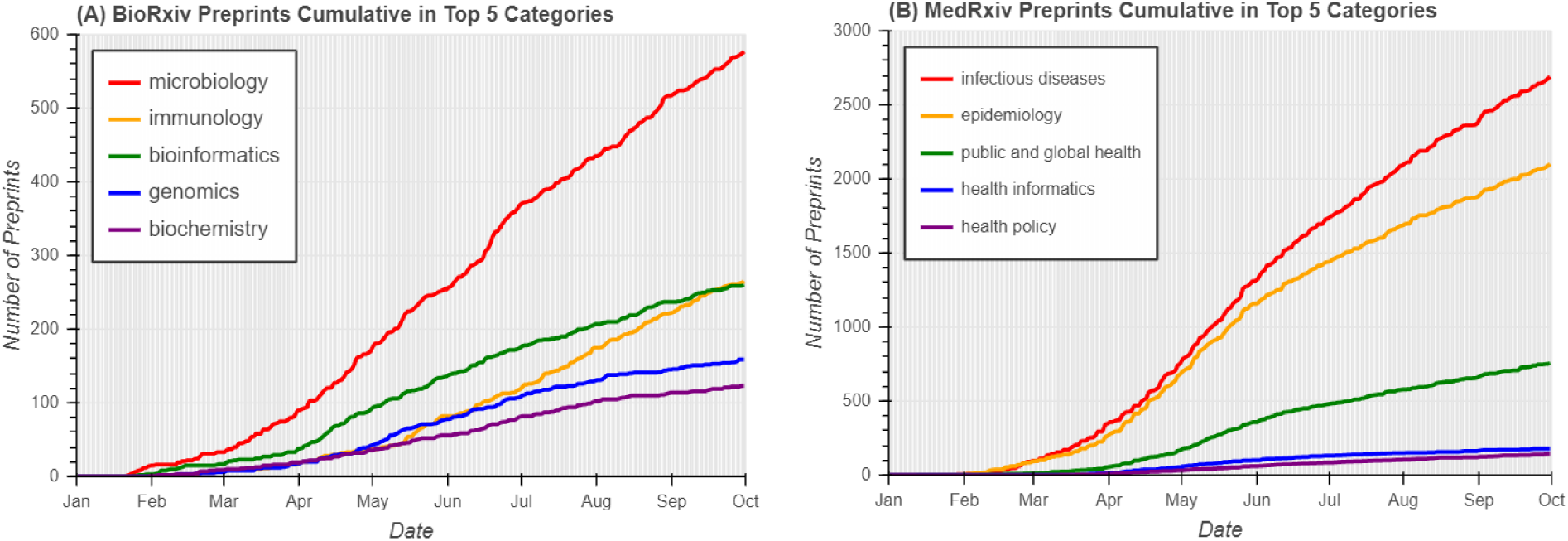
Cumulative COVID-19 preprint submissions in the top 5 categories. (A) BioRxiv and (B) medRxiv.

We also analyzed categories for bioRxiv preprints unrelated to COVID-19 (see Categories Analysis in SI) deposited into the server during Jan 1 – Sept 30, 2020. We found that the majority of preprints unrelated to COVID-19 were deposited into categories of *neuroscience*, *microbiology*, *and bioinformatics*. In fact, *neuroscience* was the leading category through Nov 2018 as was demonstrated in Blekhman’s study [51]. That same study named *bioinformatics*, *genomics*, and *evolutionary biology* as other leading categories. Interestingly, the field of *bioinformatics* remains among the leading categories for both COVID-19 related and unrelated preprints, prior to or during the coronavirus pandemic, which demonstrates the growing importance of this field for all subject areas, its interdisciplinary nature, and the familiarity of bioinformatics research community with the preprint culture.

### 3. Published preprints

Preprints deposited on a preprint server can later undergo a peer-review process to be published in refereed journals. Here, we will refer to a preprint that has a peer-reviewed journal article analogue as a published preprint. Once a preprint is published in peer-reviewed journal, preprint and article’s DOIs are permanently linked in indexing databases and on the preprint server; nevertheless, establishing this association can sometimes be challenging. For instance, the title of the research study or the authors’ list may change during the publication process, complicating auto-matching the preprint and its published version. Computing the number of preprints published as journal articles requires tedious analysis and the design of matching algorithms [59]. To ensure we find all published preprints, we combined data from various APIs (BioRxiv API, Crossref, Dimensions, Rxivist, and CORD-19) and matched publications based on both, DOIs and publication titles (see Published Collections in SI). We found that during Jan 1 – Sept 30, on average, 18% of COVID-19 preprints deposited to bioRxiv and medRxiv servers resurfaced as peer-reviewed journal articles.

As expected, the majority of published preprints were those deposited into the leading categories: *microbiology*, *immunology*, and *bioinformatic*s represented 59% of all published bioRxiv preprints, while *infectious diseases*, *epidemiology*, and *public and global health* contained 76% of all published medRxiv preprints (Fig 6). The publication rates vary across the preprint categories (Fig 7). Thus, COVID-19 preprints in bioRxiv categories of *microbiology* and *biochemistry* display the highest publication rates of 22%. The *infectious diseases* category in medRxiv contained 18% of preprints that later appeared as journal articles and it alone provided 43% of all published medRxiv preprints on COVID-19. At the same time, other leading categories, *epidemiology* and *public and global health*, both had publication rates of 13%, which is lower than an average rate for medRxiv. The lowest publication rates were observed for COVID-19 preprints deposited into the *neuroscience* category in bioRxiv and *health economics* in medRxiv. The highest publication rate of 30% was observed for *pathology* in medRxiv, although it was only 10% for bioRxiv.

**Fig 6.**
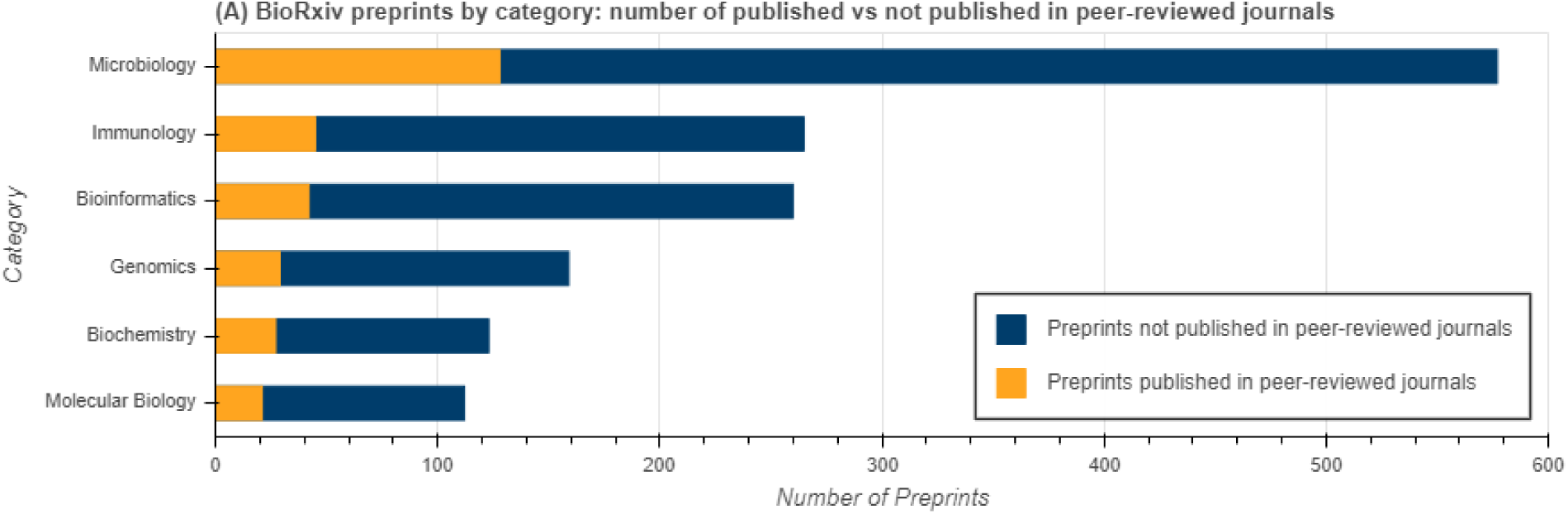
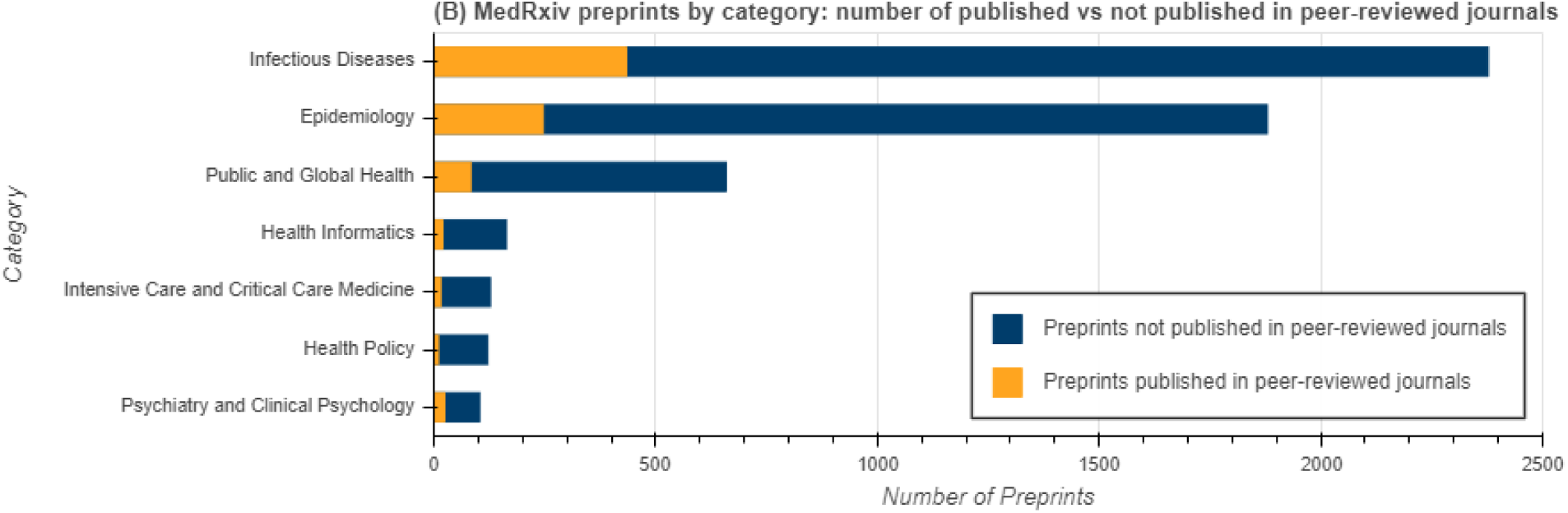
Published vs. not published COVID-19 preprints by category. (A) BioRxiv and (B) medRxiv preprints that have journal article analogues (yellow) and those that exist only as preprints (blue). The total number of preprints in the category is the sum of orange and blue bars. Only categories with at least 100 preprints are displayed for clarity purposes.

**Fig 7.**
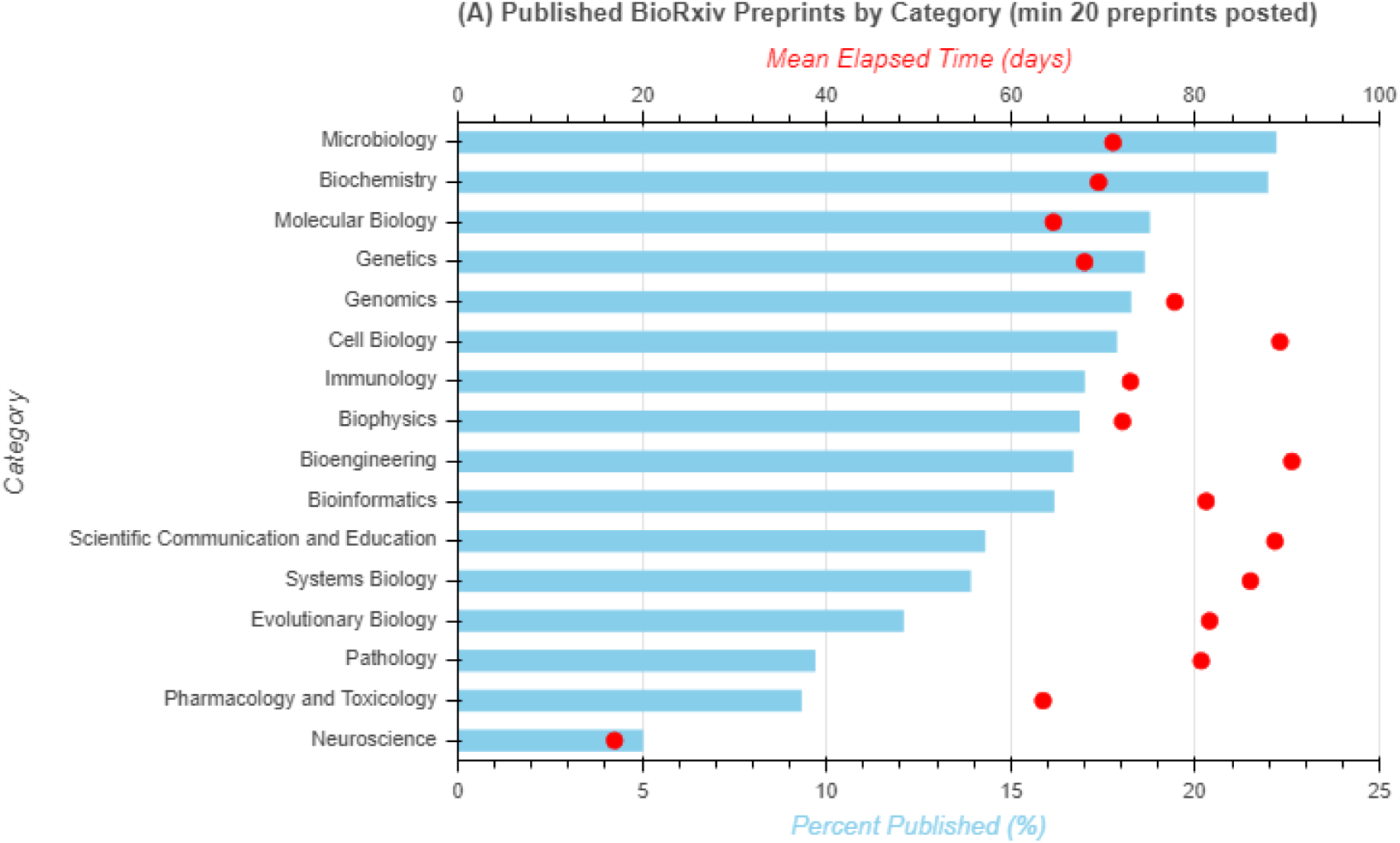
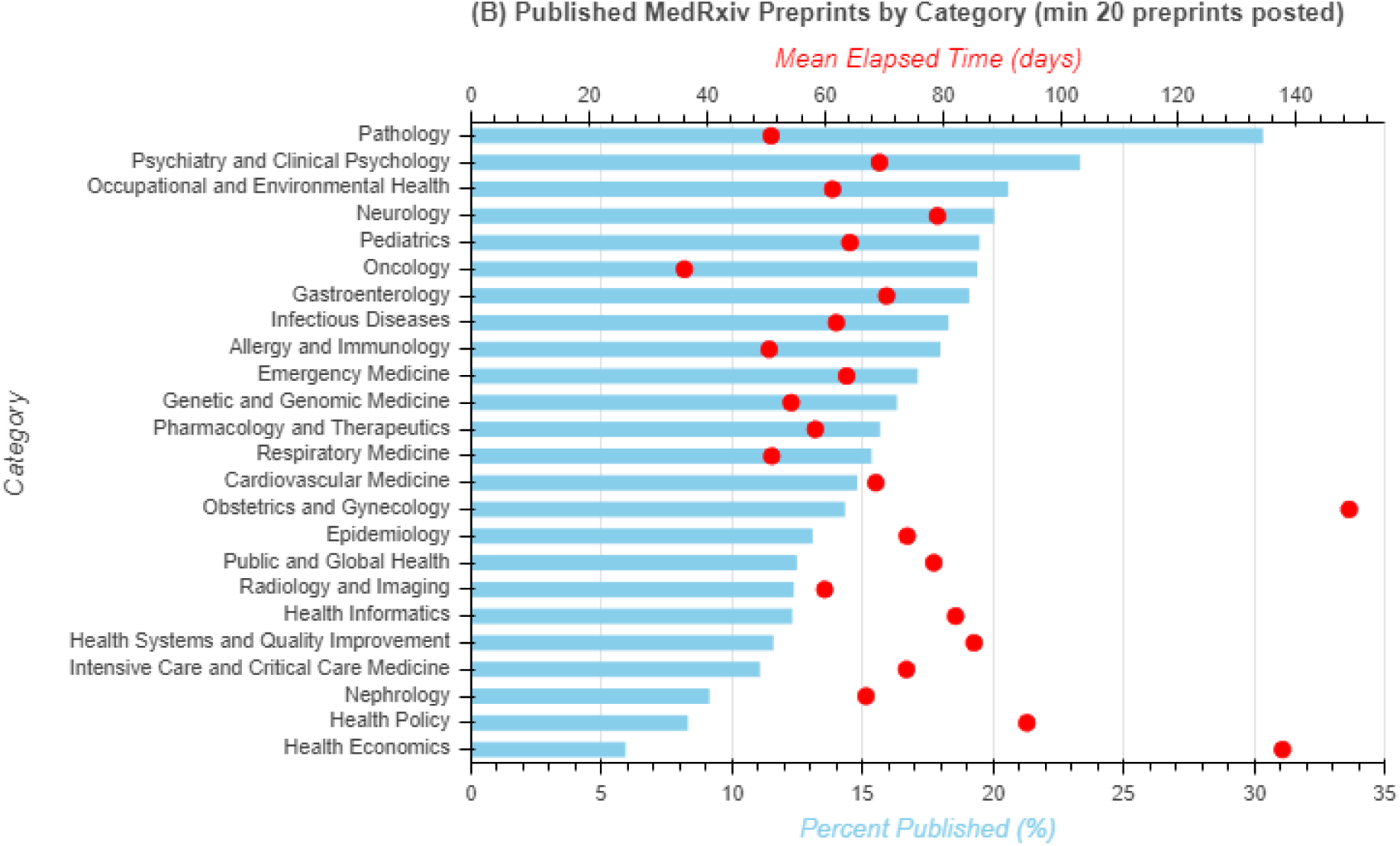
Published COVID-19 preprints (in blue) and mean elapsed time (*TΣ*, in red) by category. (A) BioRxiv and (B) medRxiv categories with at least 20 preprints published during Jan 1 – Sept 30, 2020.

The publication rates of 18% observed for COVID-19 related preprints on Sept 30, is low as compared to publication rate of 42% [51] or 70% [3] reported for pre-pandemic preprints or for previous health crises. For instance, for Ebola, 60% of the deposited preprints were published in PubMed-indexed journals, and for Zika, the corresponding amount was 48% [60]. One limitation to our study is that articles that are undergoing the peer-review or various editorial processes are invisible to us. The dependence of publication rates on the analysis timespan was noted by Blekhman *et al*. [51], who showed that publication rate for preprints was 67% during the comprehensive study that covered the period of several years (2013-2016) but only 20% during the last year of their study (2018) [51]. Similarly, our reported publication rates are higher than rates reported by two independent studies of COVID-19 preprints deposited by the end of April 2020. One study reported that 6.1% of 2,102 COVID-19 related preprints from Dimensions resurfaced as peer-review journal articles [61], while another study reported that only 4% of COVID-19 related bioRxiv and medRxiv preprints were published [62]. We further evaluated the possibility that publication rates for COVID-19 preprints deposited into medRxiv and bioRxiv servers between Jan 1 and Sept 30 were underestimated, especially considering the length of the publication process for COVID-19 related preprints (see next section). To that end, we reanalyzed publication rates on Dec 7, 2020, for COVID-19 preprints deposited between Jan 1 and Sept 30, 2020, and we found them at 34% and 29%, for bioRxiv and medRxiv preprints, respectively. Despite being higher than the publication rate of 18% derived from our data in October, reanalyzed publication rates are still low.

### 4. Publication delays for published preprints

**Scheme 1.**
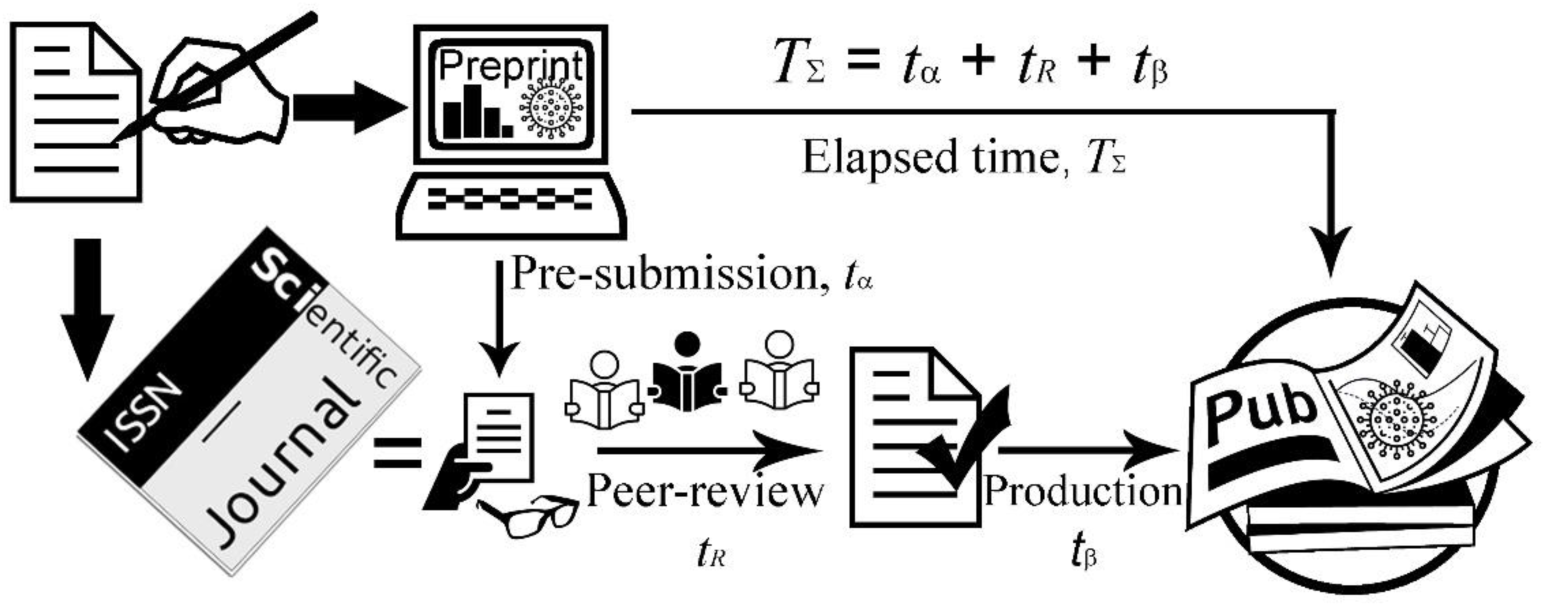
Preprint publication process [63].

We analyzed various delays involved in the publication process using the model depicted in Scheme 1. In our proposed model, the interval between preprint posting on a preprint server and online publication of the corresponding peer-reviewed journal article constitutes the *elapsed time* (*TΣ*), which can be further expressed as:

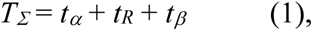

where *pre-submission time* (*tα*) is the interval between preprint posting on a preprint server and its submission to the peer-reviewed journal; *peer-review time* (*t_R_*) is the duration of the peer-review process; and *production stage time* (*tβ*) is the interval between article official acceptance statement and its publication online.

The descriptive statistics for these publication delays are summarized in Table 2 and Fig 8 and will be discussed in detail below. It is worth noting that none of the publication delays display a standard Gaussian distribution (Fig 9), thus we discuss both their medians and means.

**Fig 8.**
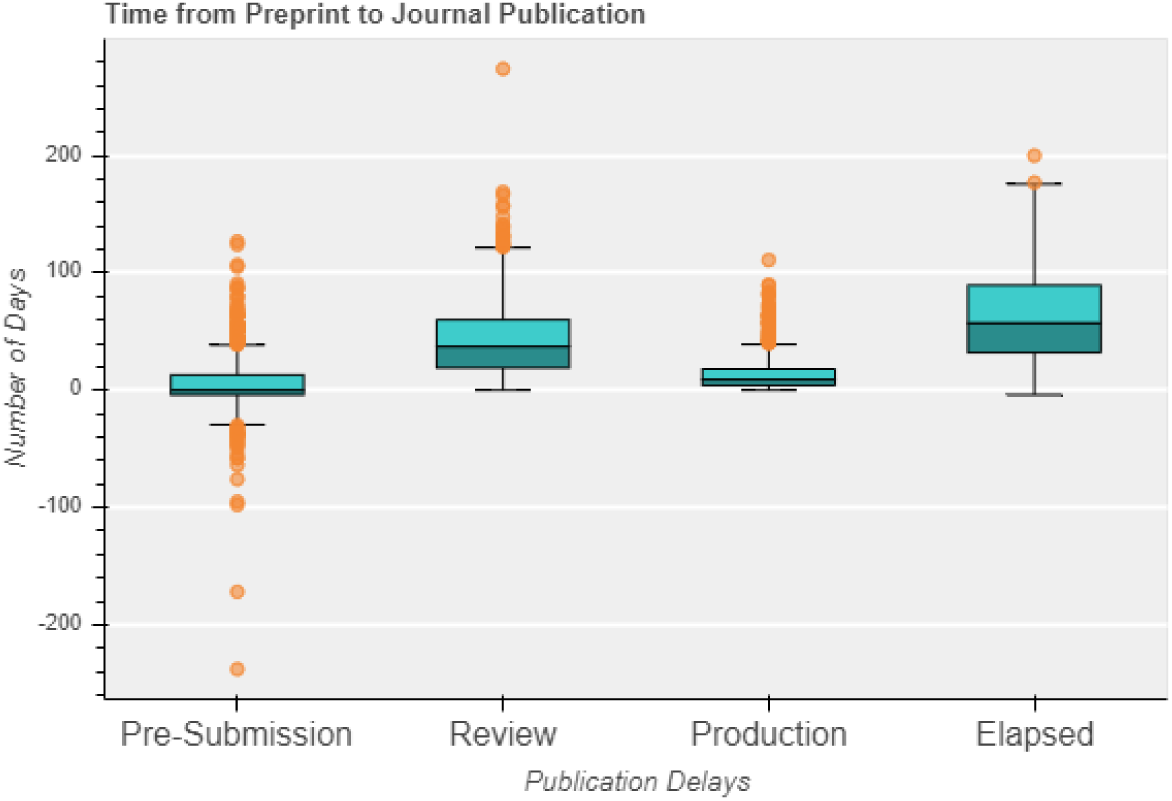
Time from preprint to journal publication. Box plot displaying publication delays for combined medRxiv and bioRxiv COVID-19 preprints deposited during Jan 1 – Sept 30, 2020.

**Fig 9.**
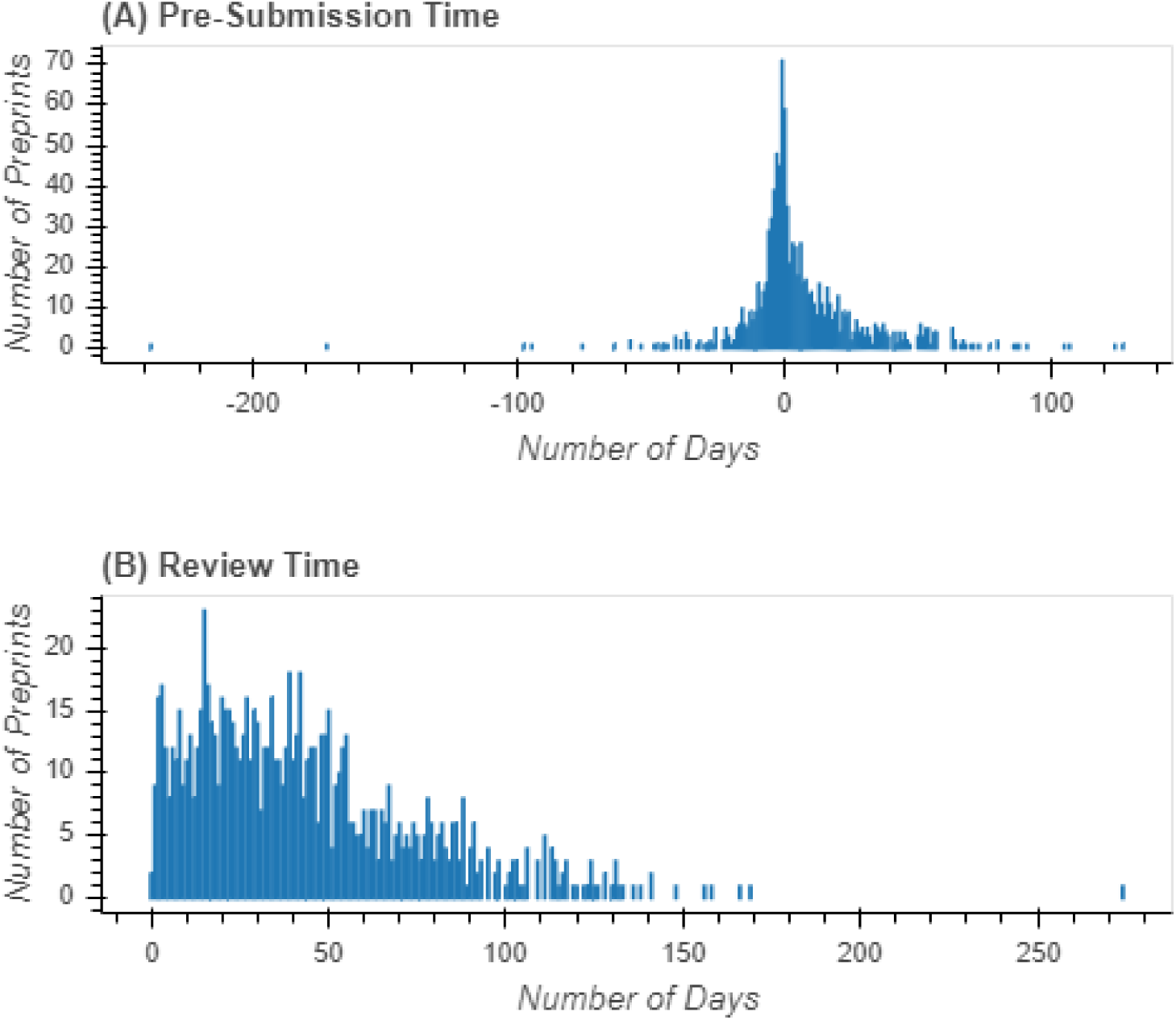
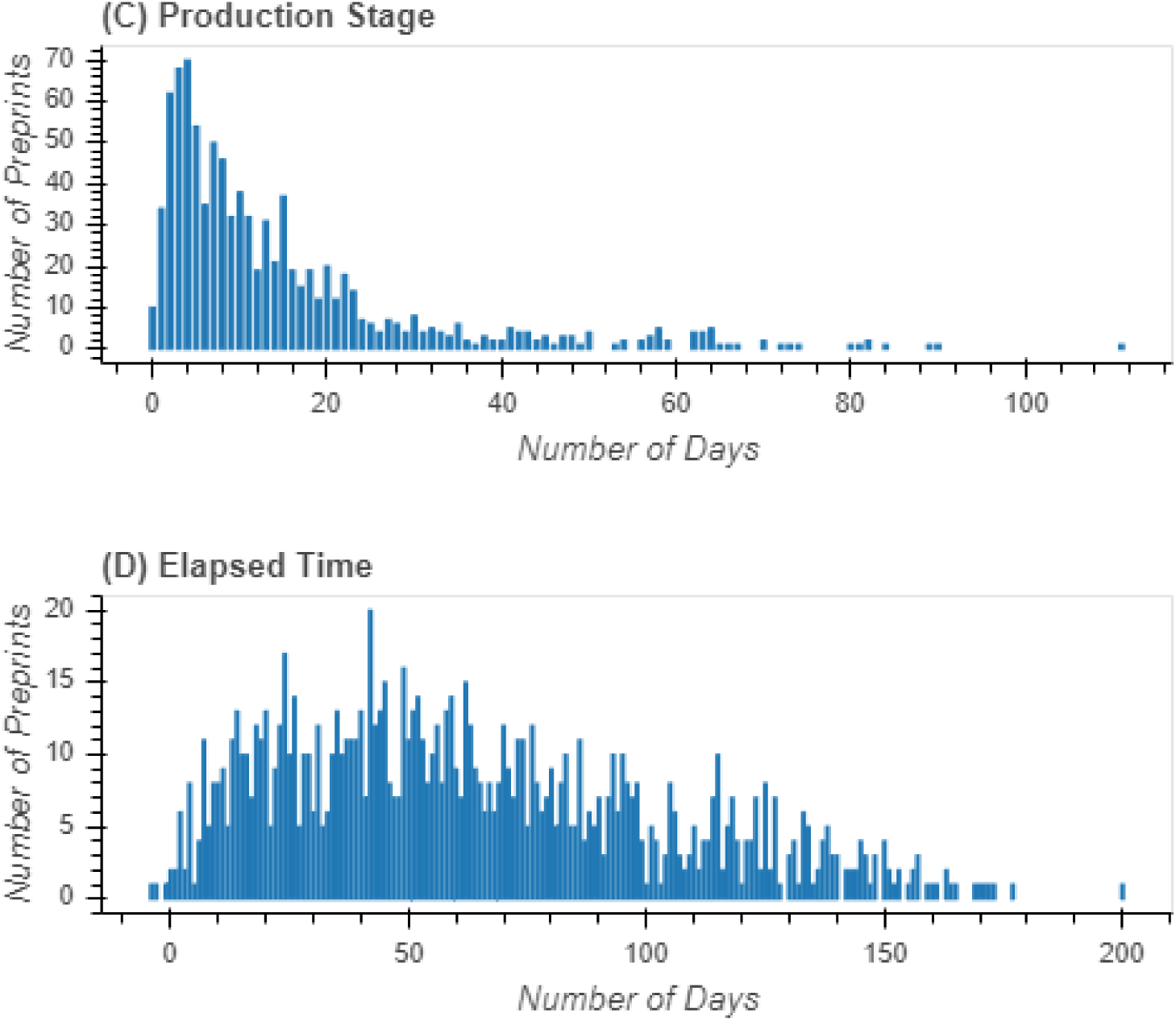
Publication delays from preprint to journal publication. Number of COVID-19 combined medRxiv and bioRxiv preprints with given number of days that correspond to various publication delays: (A) *Pre-submission time*; (B) *Review time*; (C) *Production stage time*; and (D) *Elapsed Time*.

**Table 2.** Publication delays (in days) for combined medRxiv and bioRxiv COVID-19 preprints deposited during Jan 1 – Sept 30, 2020.

| Publication delays | Symbol | Mean | SD | Median | IQR | Mode | N |
| --- | --- | --- | --- | --- | --- | --- | --- |
| *Pre-submission* | $t_{\alpha}$ | 5.6 | 23.9 | 0 | 17 | -1 | 973 |
| *Review* | $t_R$ | 43.4 | 32.4 | 37 | 41 | 15 | 950 |
| *Production* | $t_{\beta}$ | 14.6 | 15.7 | 9 | 14 | 4 | 916 |
| *Elapsed time* | $T_{\Sigma}$ | 63.4 | 39.5 | 57 | 58 | 42 | 1099 |

#### 4.1. Elapsed Time (*TΣ*) - Time from a preprint to its journal article version

For bioRxiv COVID-19 preprints, the evaluated *elapsed time* (*TΣ*) varies anywhere between 0 and 177 days, with a mean of 68.5 days (Table 3). For medRxiv COVID-19 preprints, *TΣ*varies anywhere between −4 to 200 days with the mean of 61.4 days. The difference in elapsed time between medRxiv and bioRxiv preprints is statistically significant (Table 3). The mean *TΣ*for combined bioRxiv and medRxiv COVID-19 preprints deposited between Jan 1 and Sept 30, 2020 is 63.4 days (Table 2), which is significantly faster than 166 [51] or 155 days [64] reported for bioRxiv preprints during 2013-2018; and faster than 150 days reported for Zika or Ebola preprints [60] (Table 4). Curiously, our mean *TΣ*is significantly longer than 22.5 days reported by Coates *et al*. for bioRxiv and medRxiv preprints published by the end of April 2020 [62], although their data may be unintentionally biased towards low mean *TΣ*values considering only 4% of preprints were published at that time.

**Table 3.**
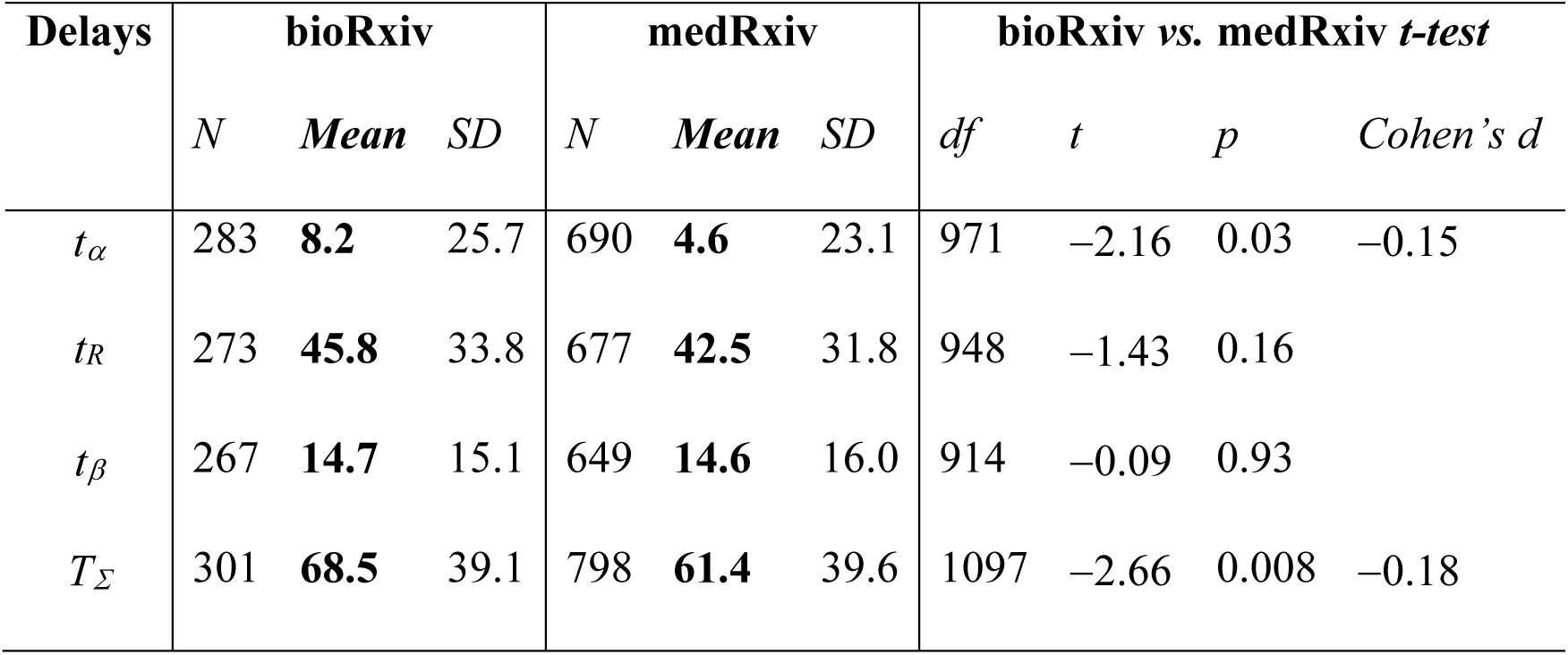
Descriptive statistics for publication delays (in days) for bioRxiv and medRxiv, as well as Student’s t-test to evaluate their discrepancy.

| Delays | bioRxiv |  |  | medRxiv |  |  | bioRxiv vs. medRxiv *t*-test |  |  |  |
| --- | --- | --- | --- | --- | --- | --- | --- | --- | --- | --- |
|  | *N* | *Mean* | *SD* | *N* | *Mean* | *SD* | *df* | *t* | *p* | *Cohen's d* |
| $t_{\alpha}$ | 283 | **8.2** | 25.7 | 690 | **4.6** | 23.1 | 971 | -2.16 | 0.03 | -0.15 |
| $t_R$ | 273 | **45.8** | 33.8 | 677 | **42.5** | 31.8 | 948 | -1.43 | 0.16 | |
| $t_{\beta}$ | 267 | **14.7** | 15.1 | 649 | **14.6** | 16.0 | 914 | -0.09 | 0.93 | |
| $T_{\Sigma}$ | 301 | **68.5** | 39.1 | 798 | **61.4** | 39.6 | 1097 | -2.66 | 0.008 | -0.18 |

**Table 4.** Elapsed time (TΣ, in days) for COVID-19 preprints as compared to previous studies (df = 1098).

| $T_{\Sigma}$ | Retrieval of article publication dates | *Student's t*-test | | |
| --- | --- | --- | --- | --- |
| [Ref] |  | *t* | *p* | *Cohen's d* |
| 166 | BioRxiv preprints (2013-2018). Date corresponds to | -86.06 | < 0.001 | -2.60 |
| [51] | Crossref's "published-online" date or, when not available, "published-print" date. |  |  |  |
| 155 | BioRxiv preprints (2013-2017). Date corresponds to | -76.84 | < 0.001 | -2.32 |
| [64] | Crossref's "created-date". |  |  |  |
| 150 | Zika (2015-2017) and Ebola (2014-2016) related | -72.65 | < 0.001 | -2.19 |
| [60] | preprints. Date corresponds to "pub_date" in PubMed. |  |  |  |
| 22.5 | COVID-19 medRxiv and bioRxiv preprints (Jan– | 34.24 | < 0.001 | 1.03 |
| [62] | April, 2020). Dates retrieval method is not specified. |  |  |  |

We also explored whether *TΣ*can explain the different publication rates for preprint categories. For example, medRxiv preprints in *health policy* (*M* = 94.3, *SD* = 41.1, *N* = 10) and *health informatics* (*M* = 82.2, *SD* = 41.8, *N* = 20) have long mean *TΣ*and display low publication rates, while preprints in *pathology* display the shortest *TΣ*(*M* = 50.9, *SD* = 33.0, *N* = 10) and, in fact, have high publication rate (Fig 7 and Categories Analysis in SI). On the contrary, bioRxiv preprints in *cell biology* display the longest *TΣ*(*M* = 89.2, *SD* = 46.4, *N* = 10) but not the lowest publication rate. Pearson’s analysis showed that *elapsed time* and publication rate do not correlate for bioRxiv preprints and only slightly correlate for medRxiv preprints (for medRxiv: *r*(41) = −0.39*, p* = 0.011; for bioRxiv: *r*(19) = −0.13*, p* = 0.581). As publication rates settle, we plan to re-analyze whether the disciplinary publication trends are affected by the duration of the publication process.

#### 4.2. Review time (*tR*) and production stage time (*tβ*)

While the *elapsed time* describes how quickly it takes for a preprint to resurface as a journal article, it is unable to distinguish whether the successful expediting of a publication process is a result of the new policies implemented by journal publishers or the old practices of posting a preprint on a preprint server shortly after or even prior to its submission to the peer-reviewed journal. In our quest to explain the expedited publication times, we analyzed *review time* (*t*_R_) and the *production stage period* (*tβ*) (Fig 10).

**Fig 10.**
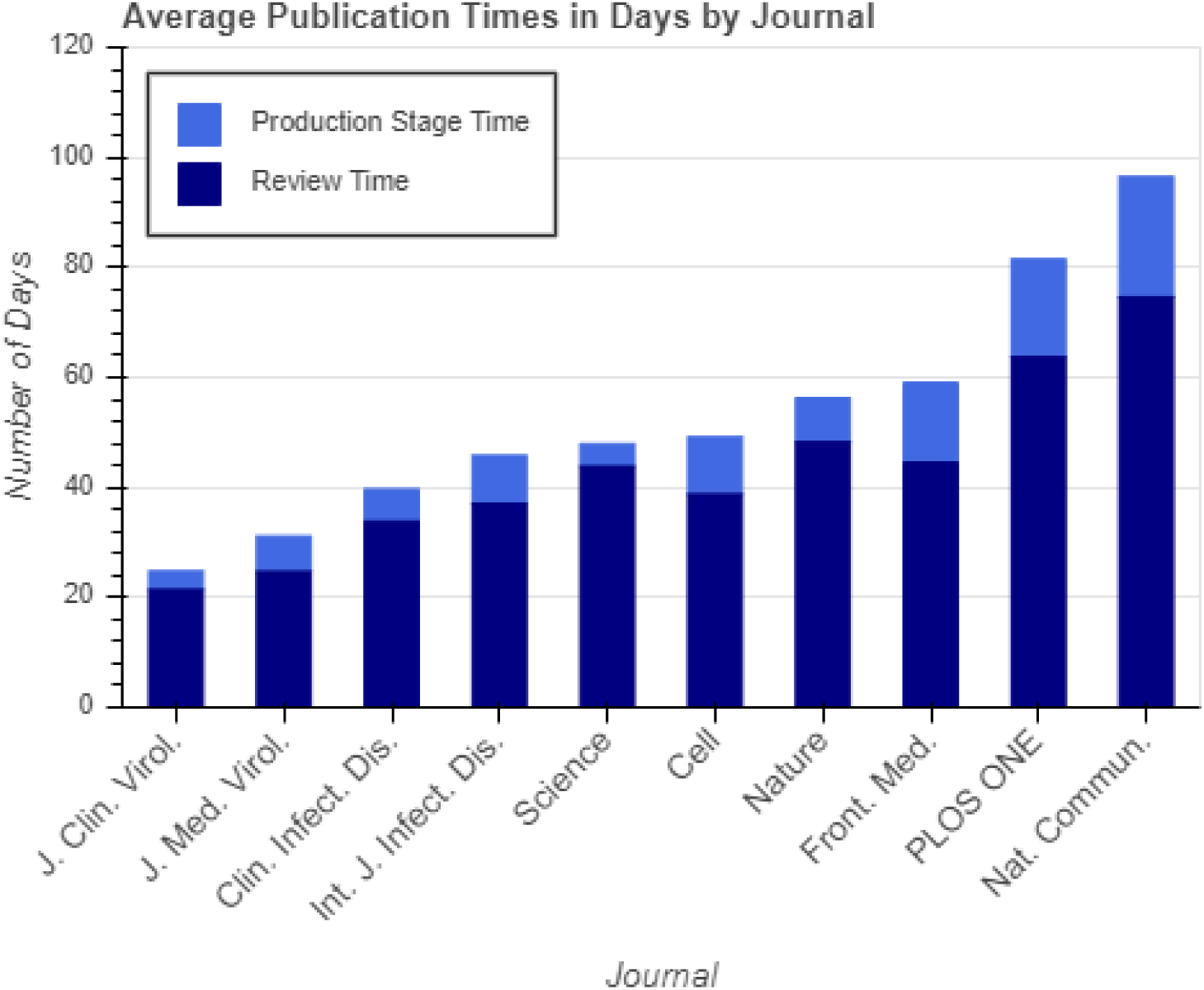
Mean publication delays. *Review time* (*t*_R_, dark blue bar) and *production stage time* (*tβ*, light blue bar) are displayed in days for the top 10 journals publishing COVID-19 medRxiv and bioRxiv preprints. The entire two-colored bar represents the total publication time.

We found that a mean *review time* (*t*_R_) for COVID-19 related bioRxiv and medRxiv preprints is 43.4 days (Table 2), which is significantly shorter (*t*(949) = −53.85, *p* < 0.001, *Cohen’s d* = −1.75) than a standard *review time* of 100 days [65]. The difference in the mean *t*_R_ for COVID-19 related medRxiv and bioRxiv preprints is not statistically significant (Table 3). Our data showed that *t_R_* for COVID-19 articles has significantly decreased (*t*(949) = −92.86, *p* < 0.001, *Cohen’s d* = −3.01), by about 70%, during the pandemic, as compared to the earlier analysis by Björk and Solomon, who in 2013 reported that biomedicine journals exercised a mean *t*_R_ of 141 days [66]. We found that *t*_R_ for COVID-19 articles associated with preprints was significantly shorter than 51.0 days (*t*(949) = −7.23, *p* < 0.001, *Cohen’s d* = −0.24) reported for all coronavirus articles [61]; and significantly shorter than 84 days (*t*(949) = −38.63, *p* < 0.001, *Cohen’s d* = −1.25) reported for articles published on topics other than COVID-19 during 2020 [67]. Although both studies describe the period of early pandemic, since January and until the end of April, the former study reports a mean review time of 51.0 days [61], while the latter a median *review time* of 6 days [67]. This discrepancy in early data is likely due to a sever skew in frequency distribution for *t*_R_.

The major advance in speeding up the peer-review process was observed for *PLOS ONE*. For COVID-19 related articles published in *PLOS ONE* between Jan 1 and Oct 23, 2020, the peer-review took on average 63.7 days. This is significantly shorter than 125 days in 2016 [68] (*t*(59) = −−15.13, *p* < 0.001, *Cohen’s d* = −1.95) and 126.6 days in April 2020 [61] (*t*(59) = −15.52, *p* < 0.001, *Cohen’s d* = −2.00). Of the top ten journals that published medRxiv and bioRxiv COVID-19 preprints, *Journal of Clinical Virology* had the shortest mean review time of 21.4 days and *Nature Communications* had the longest mean review time of 74.6 days (Table 5, Fig 10).

**Table 5.** Descriptive statistics for publication delays (in days) for the top 10 journals that published medRxiv and bioRxiv COVID-19 preprints.

| *Journal* | N | Review |  | Production |  | Publication |  | Elapsed Time |  |  |
| --- | --- | --- | --- | --- | --- | --- | --- | --- | --- | --- |
| | | $(t_R)$ | | $(t_\beta)$ | | $(t_R + t_\beta)$ | | $(T_\Sigma)$ | | |
|  |  | Mean | SD | Mean | SD | Mean | SD | Mean | SD | N |
| *J. Clin. Virol.* | 15 | **21.4** | 41.7 | **3.3** | 2.3 | **24.7** | 41.2 | **26.2** | 29.7 | 18 |
| *J. Med. Virol.* | 26 | **24.7** | 21.7 | **6.5** | 4.5 | **31.1** | 22.3 | **38.3** | 28.8 | 27 |
| *Clin. Infect. Dis.* | 18 | **33.8** | 13.9 | **6.8** | 13.0 | **39.7\*** | 20.3* | **46.8** | 33.5 | 32 |
| *Int. J. Infect. Dis.* | 23 | **37.0** | 20.3 | **8.7** | 9.1 | **45.8** | 19.7 | **53.7** | 31.1 | 24 |
| *Science* | 21 | **43.8** | 37.4 | **4.1** | 2.3 | **47.9** | 37.8 | **51.2** | 41.8 | 25 |
| *Cell* | 15 | **38.7** | 21.4 | **10.4** | 7.8 | **49.1** | 16.4 | **43.5** | 16.4 | 15 |
| *Nature* | 13 | **48.2** | 27.0 | **8.0** | 2.0 | **56.3** | 28.3 | **52.3** | 28.4 | 14 |
| *Front. Med.* | 15 | **44.5** | 20.5 | **14.5** | 4.7 | **59.1** | 21.1 | **71.8** | 39.0 | 18 |
| *PLOS ONE* | 60 | **63.7** | 31.4 | **17.8** | 7.3 | **81.5** | 32.5 | **89.4** | 36.2 | 83 |
| *Nat. Commun.* | 19 | **74.6** | 58.9 | **21.9** | 9.8 | **96.5** | 66.4 | **102.7** | 30.9 | 23 |

\* - based on 26 publications.

The mean *production stage time* (*tβ*) for COVID-19 related bioRxiv and medRxiv preprints is 14.6 days, about one third of the average *t*_R_ found above for the same set of articles (Table 2). The difference in *tβ*for medRxiv and bioRxiv preprints on COVID-19 is not significant (Table 3). As compared to the *tβ*of 147 days reported by Björk and Solomon in 2013 [66], *tβ*reduced by 90% for COVID-19 articles associated with preprints; this difference being statistically significant (*t*(915) = −254.89, *p* < 0.001, *Cohen’s d* = −8.42). However, a study early in the pandemic reported a significantly shorter *tβ*= 9.3 days [61] (*t*(915) = 10.17, *p* < 0.001, *Cohen’s d* = 0.34), although this value was obtained for all published coronavirus articles, not just those associated with preprints. Of the top ten journals that published COVID-19 bioRxiv and medRxiv preprints, *Journal of Clinical Virology* has the shortest *tβ*of 3.3 days and *Nature Communications* has the longest *tβ*of 21.9 days (Table 5, Fig 10).

#### 4.3. Pre-submission time (*tα*)

We found that an average *pre-submission time* (*tα*) for COVID-19 related preprints is 5.6 days (Table 2), a positive value implying that, on average, authors posted their manuscript to the preprint server before advancing their preprints to journal publishers (Fig 11). Authors of bioRxiv COVID-19 preprints waited longer than authors of medRxiv COVID-19 preprints; this difference being statistically significant (Table 3). The distribution of *tα*frequencies indicates a median at 0 days (Table 2, Fig 9). A more detailed analysis showed that 44% of the COVID-19 preprints were deposited to bioRxiv or medRxiv servers after being submitted to the journal (negative *tα*) and only 28% of preprints were posted more than 10 days before they were submitted to the journal where they were published. Our results mirror earlier findings by Anderson [69], who reported those values as 57% and 29%, respectively, for papers that had preprint analogues and were published in *Nature* journals between 2013 and 2018. This authors’ behavior was also observed for arXiv preprints [70] and explained by a widespread fear for “getting scooped” when making details of research publicly available. We thus confirm that the previously reported trends are restated for COVID-19 preprints deposited between Jan 1 and Sept 30, 2020. We believe that during the pandemic, bioRxiv and medRxiv preprint servers were not used to gather a community feedback because authors primarily pursued rapid and open dissemination of critical research.

**Fig 11.**
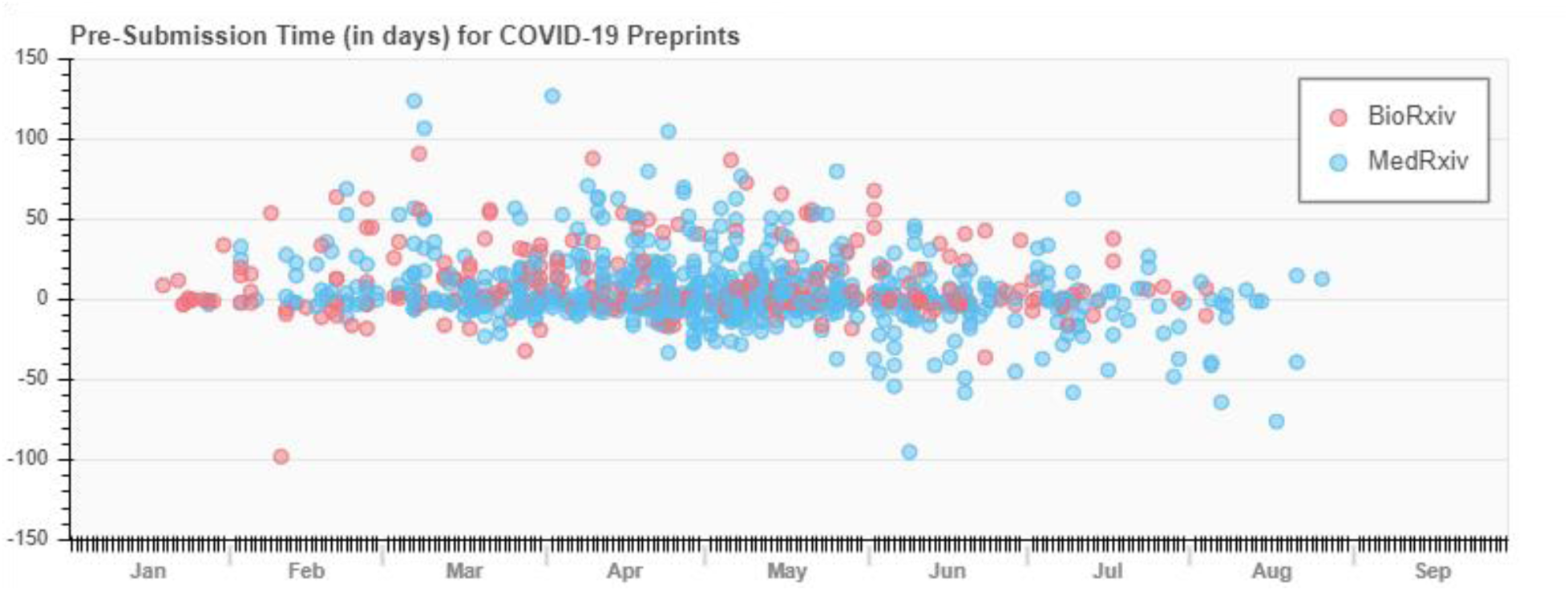
*Pre-submission time* for COVID-19 published preprints. The *tα*, in days, is plotted for bioRxiv (red) and medRxiv (blue) preprints deposited during Jan 1 – Sept 30, 2020. The 0 date is the date the preprint was submitted to the peer-reviewed journal and a positive *tα*indicates that the preprint was deposited before being submitted to the journal.

### 5. Journals

#### 5.1. Top journals by number of submissions

By Oct 23, 2020, 1,359 preprints from both, medRxiv and biorxiv appeared as peer-reviewed journal articles in 525 different academic journals, with only 25 journals (4.8%) publishing at least ten preprints (see Leading Journals in SI). The top three journals publishing the majority of bioRxiv preprints were *Science*, *Nature*, and *Nature Communications*. The top three journals publishing the majority of medRxiv preprints and, in fact, the majority of combined medRxiv and bioRxiv COVID-19 related preprints were *PLOS ONE*, *Clinical Infectious Diseases*, and *Journal of Medical Virology*.

Previous analysis of bioRxiv preprints showed that *Scientific Reports*, *eLife*, and *PLOS ONE* published the majority of server’s preprints by the end of 2018 [51]. While *PLOS ONE* remains a leader in publishing COVID-19 related preprints, the majority of them (91.6%) are from medRxiv. *Scientific Reports* during this period published only 0.4% of COVID-19 articles, of which 20.7% were originally preprints and 75% of which were specifically from bioRxiv. In comparison, two studies early in the pandemic reported that the majority of preprints were published by *Nature*, *Cell*, and *Science* (for COVID-19 preprints indexed in Dimensions) [61] and by *Journal of Medical Virology* (for COVID-19 bioRxiv and medRxiv preprints) [37].

#### 5.2. Elapsed time (*TΣ*) by journals

We reasoned that variations in the set of journals publishing the majority of preprints could be explained with difference in elapsed times for each journal (Fig 12). Indeed, in *post hoc* Bonferroni-corrected pairwise comparison, we found that the *Journal of Medical Virology*, *Nature*, *Cell*, and *Science* had significantly shorter elapsed time (*TΣ*) as compared to *PLOS ONE* (*p* < .001 − 0.03). A one-way ANOVA test showed that there was a significant main effect (*F*(5, 175) = 19.33, *p* < .001, *η*_p_^2^ = 0.36). The other four journals did not differ between each other. This explains why early in the pandemic, *Journal of Medical Virology*, *Nature*, *Cell*, and *Science* incorporated more published preprints than *PLOS ONE*. As can be seen from Fig 12, it took longest for COVID-19 preprints to be published in *Nature Communications* or *PLOS ONE*, and quickest - in *Journal of Clinical Virology* (Table 5).

**Fig 12.**
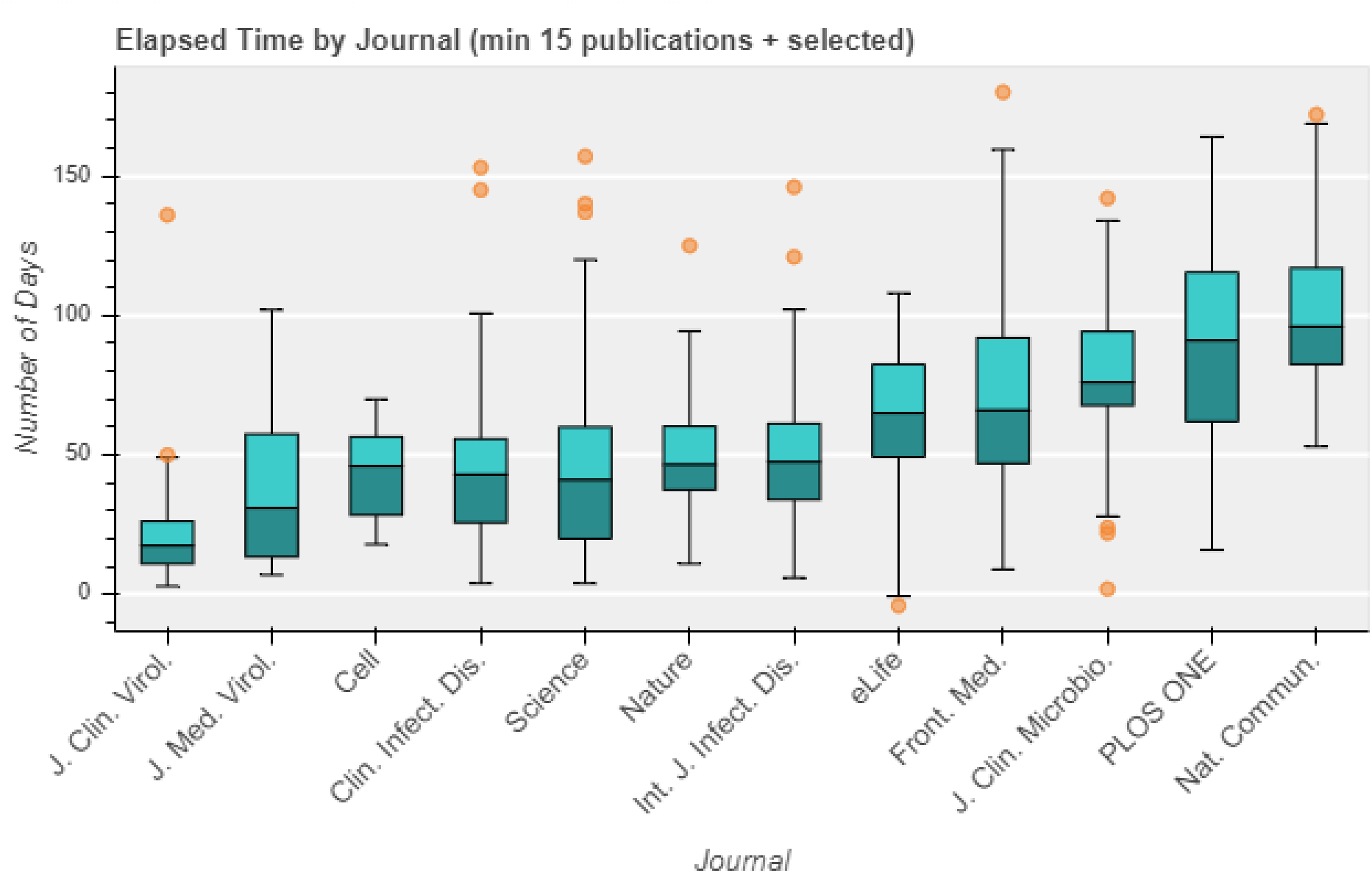
E*lapsed time* by journal. Box plot depicts quartiles (0.0, 0.25, 0.5, 0.75, 1.0) and outlier data for *TΣ*of combined COVID-19 related medRxiv and bioRxiv preprints published as journal articles in selected journals.

#### 5.3. Alternative ranking for top journals

In the previous section, we discussed journals that published the majority of bioRxiv and medRxiv preprints based on the number of preprints. By this metrics, mega-journals that publish a large number of articles will systematically appear as leaders in publishing preprints. An alternative way to allocate leading journals is to evaluate what fraction of journal’s publications come from preprints. For this, we analyzed the percentage of published COVID-19 related preprints with respect to all journal and review articles in a particular journal, as indexed in PubMed between Jan 1 and Sept 30, 2020. We found that COVID-19 preprints published in the mega-journal *PLOS ONE* constituted only 0.6% of its total publication volume (see Leading Journals in SI). However, if we only compare to the research and review articles on COVID-19, we find that 20.8% of *PLOS ONE* COVID-19 articles are former preprints. A similar scenario is observed for other multidisciplinary journals, like *Science* or *Nature*. For example, 34.8% of COVID-19 articles in *Nature Communications* come from preprints but they constitute only 0.5% of the total publication volume for this journal. The COVID-19 portfolio of *BMC Medicine* and *Journal of Clinical Microbiology* incorporated 36.4% and 33.3%, respectively, of former preprints. These percentages become 5.3% and 5.7% when we consider what fraction of all articles in *BMC Medicine* and *Journal of Clinical Microbiology*, respectively, are COVID-19 former preprints. As compared to *Nature Communications* and *PLOS ONE*, these percentages are higher, indicating that *BMC Medicine* and *Journal of Clinical Microbiology* have disciplinary scope that is adequate for coronavirus research and is more specialized than that of *Nature Communications* or *PLOS ONE*. Our analysis of COVID-19 papers associated with preprints shows that *Emerging Microbes & Infections* published 68.8% of COVID-19 related preprints, which constituted 17.7% of this journal’s COVID-19 articles. *Emerging Microbes & Infections* is an open access peer reviewed journal from Taylor & Francis, which scope, as indexed by Scopus [54], is *parasitology*, *infectious diseases*, *microbiology*, *virology*, *immunology*, *drug discovery*, and *epidemiology*, categories that are all very relevant to coronavirus research.

#### 5.4. Categories of preprints and corresponding journals

To analyze the scope of journals that published COVID-19 preprints, we plotted preprint subject categories *vs*. journal’s scope categories, as defined by Scopus [54], for all published preprints and their article analogues (Fig 13). We found that the majority of COVID-19 preprints in both medRxiv and bioRxiv were published in journals whose scope is *general biochemistry*, *genetics*, and *molecular biology*. Additionally, *microbiology* preprints from bioRxiv were published in journals specialized in *microbiology, infectious diseases*, and *virology*. The latter category is currently absent in either bioRxiv or medRxiv platforms but is listed among Scopus categories. The majority of medRxiv preprints were published in journals whose scope is *general medicine*. Preprints in *infectious diseases* and *epidemiology* were published in journals whose scope is *infectious diseases* and *microbiology (medical)*. For both, medRxiv and bioRxiv, the COVID-19 preprints in leading categories also resurfaced as journal articles in *multidisciplinary* journals, of which *PLOS ONE*, *Science*, and *Nature Communication* published the majority of them (see Leading Journals in SI).

**Fig 13.**
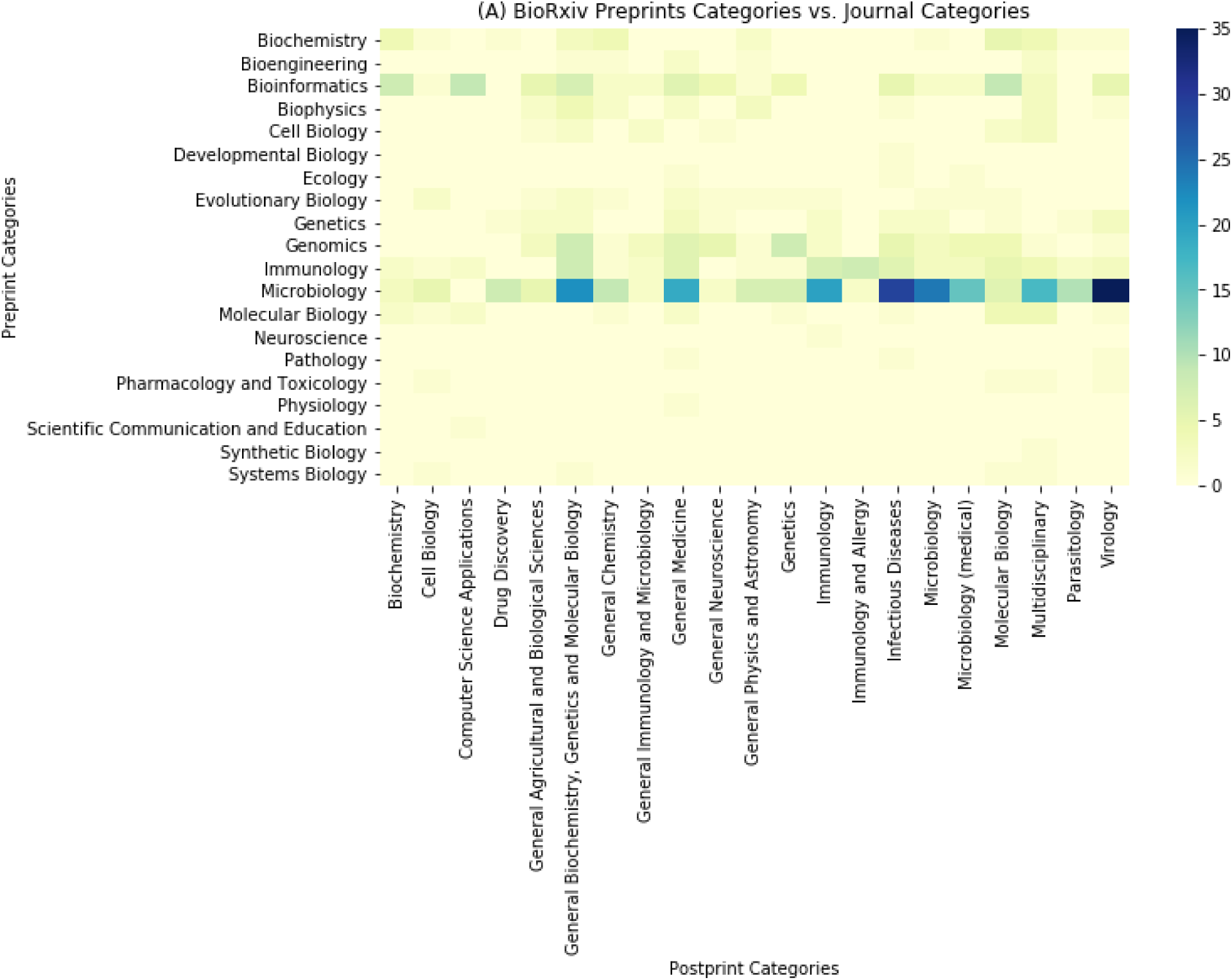
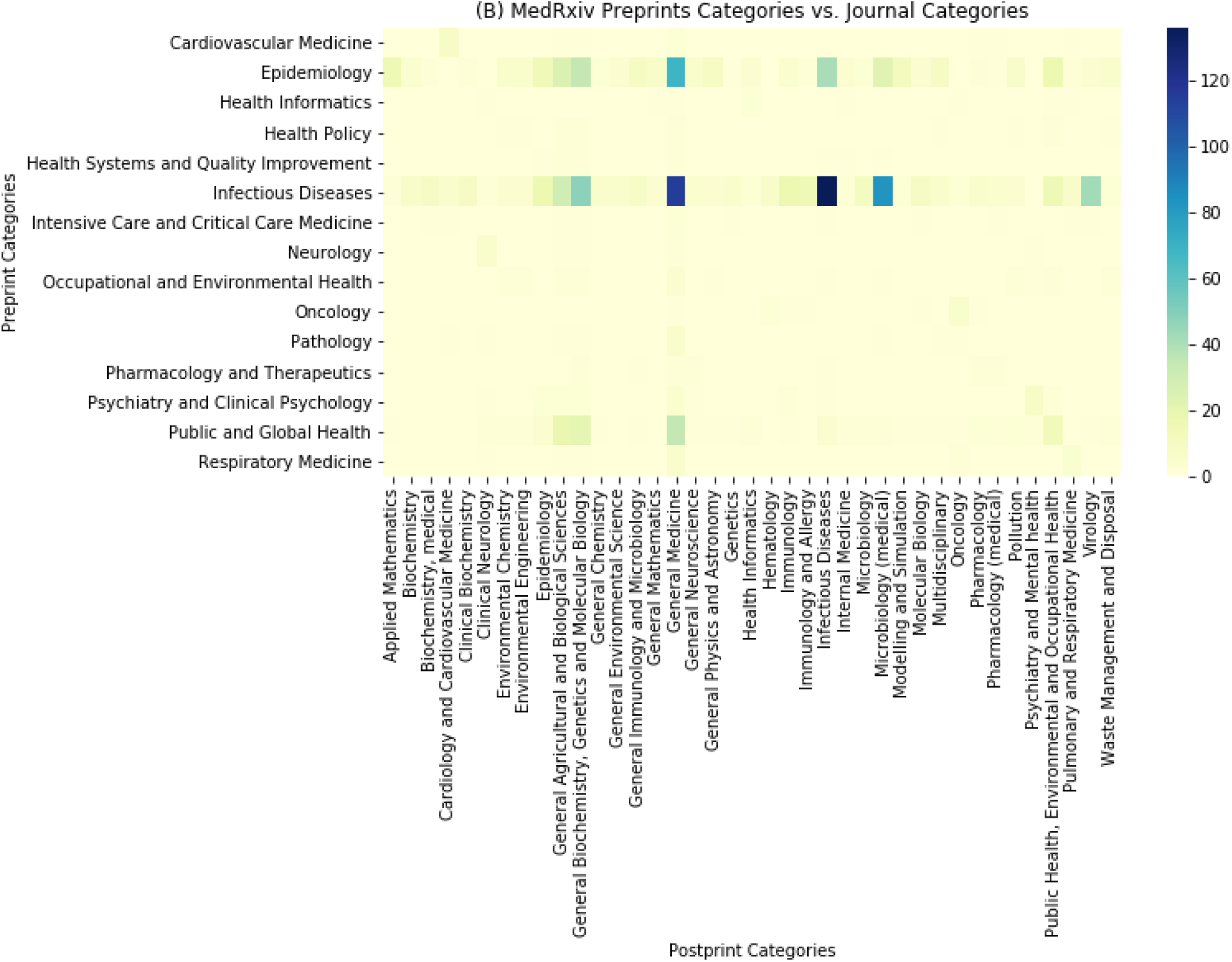
Preprint *vs.* journal categories for published COVID-19 preprints. (A) BioRxiv and (B) medRxiv preprint categories vs. journal categories of corresponding journal articles. A preprint category may be associated with many journal categories as journal scope is usually represented by multiple categories. Categories that appear less than 10 times are removed for clarity purposes.

#### 5.5. Publishers

The top two publishers for COVID-19 bioRxiv and medRxiv preprints were Elsevier and Springer. By Sept 30, they published 40% of all COVID-19 preprints deposited into bioRxiv and medRxiv during Jan 1 – Sept 30, although medRxiv preprints constituted the majority of their publications (73% in Elsevier and 65% in Springer). Elsevier journals that published the majority of COVID-19 preprints were *International Journal of Infectious Diseases*, *Journal of Clinical Virology*, and *Cell*. Springer journals that published the majority of COVID-19 preprints were *Nature Communications*, *Nature*, and *Scientific Reports*. The third major publisher for COVID-19 preprints was Oxford University Press, which journals like *Clinical Infectious Diseases*, *The Journal of Infectious Diseases*, *Journal of Travel Medicine*, and *American Journal of Clinical Pathology* published 7% of bioRxiv and medRxiv preprints on COVID-19 by Sept 30, 2020. The third major publisher for COVID-19 bioRxiv preprints was the *American Society for Microbiology* that alone published 7% of all COVID-19 preprints deposited into bioRxiv by Sept 30, 2020.

### 6. Impact of preprints

Previous studies of bioRxiv preprints deposited during 2013-2017 [64], 2014-2016 [71], and 2015-2018 [18], all ascertain an altmetric advantage of articles that were associated with preprints over articles that did not come from preprints. A notion of altmetric advantage was derived from higher Altmetric Attention Scores, which summarize various mentions of the article in the public policy documents, Wikipedia articles, mainstream news, blogs, and various social platforms [72]. For example, both bioRxiv and medRxiv preprints are widely shared on Twitter and even prior to pandemic, a single preprint was reported to generate on average 13-18 tweets [69].

To assess the visibility of COVID-19 preprints, we compared the Altmetric Attention Scores of COVID-19 related articles that had associated preprints to those that did not, and to articles unrelated to COVID-19 that were published between Jan 1 and Nov 19, 2020 (Fig 14, Table 6). We also stratified our results by journal to eliminate a potential effect of a journal’s impact factor (IF) or other journal-specific variables. For the top ten journals that published the majority of COVID-19 preprints, we found that Altmetric Attention Scores for articles that had associated preprints were slightly higher on average but not significantly different from articles that did not have associated preprints (Table 6). As expected, all COVID-19 related articles had an altmetric advantage over non-COVID-19 related publications in the same journals. For example, in *Nature*, COVID-19 related articles published between Jan 1 and Nov 19, 2020 had on average an altmetric score of 1077.8 (*SD* = 1467.8, *N* = 253). This is significantly higher (*t*(263.3) = −7.76, *p* < 0.001, *Cohen’s d* = −0.89) than an average altmetric score of 354.3 (*SD* = 722.3, *N* = 2766) for all other articles in the same journal published during the same period. Similar analysis of all ten journals resulted in average altmetric score of 332.8 for COVID-19 related works *vs*. 91.0 for non-COVID-19 publications, this difference being statistically significant (Table 6). As seen from Fig 14, altmetric scores vary for different journals. In fact, we found a strong Pearson’s correlation between the Altmetric Attention Scores for journal articles and journals’ impact factors (for COVID-19 articles that were deposited as preprints, *r*(8) = 0.99*, p* < 0.001).

**Fig 14.**
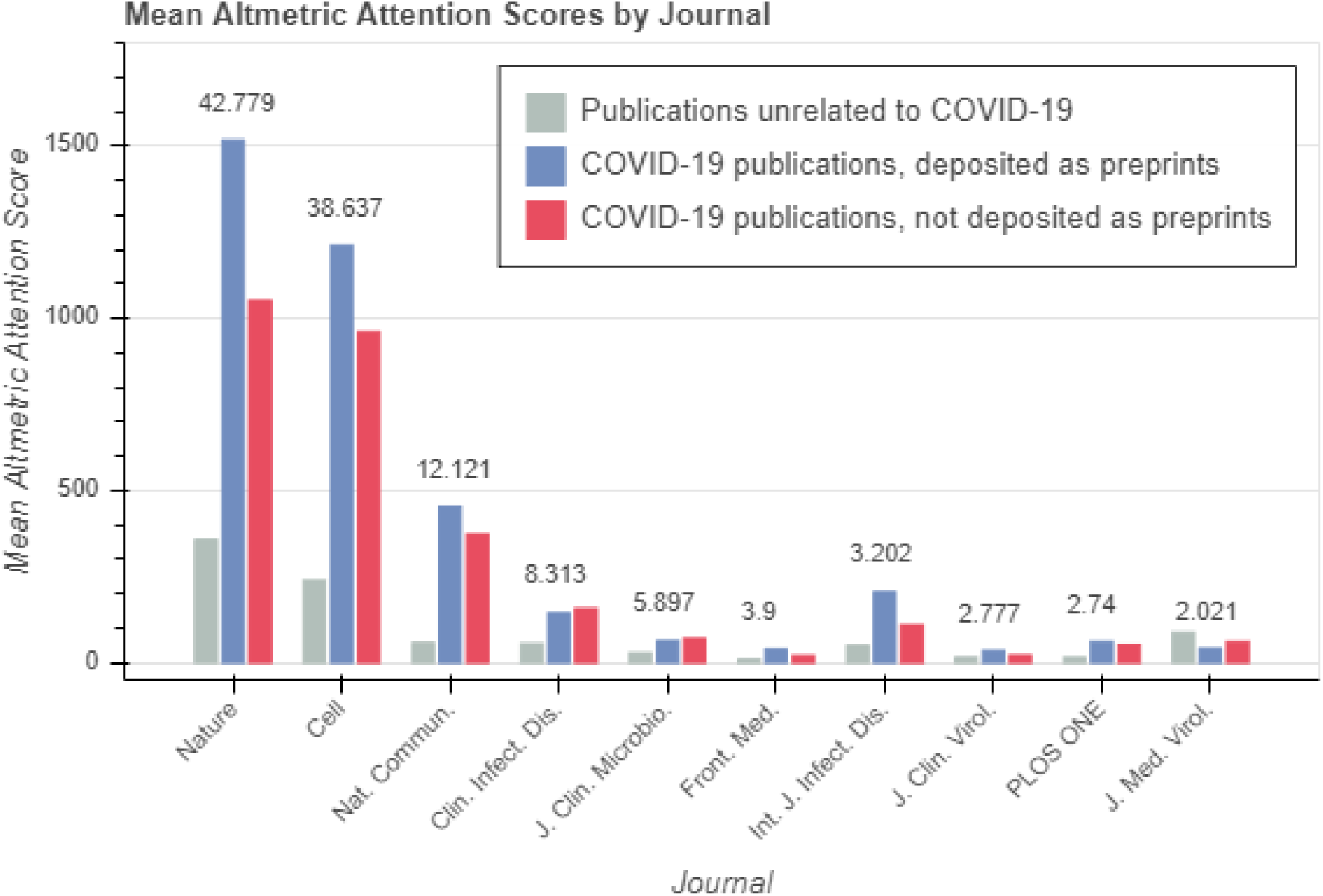
Mean Altmetric Attention Scores by journal. For COVID-19 related articles that were deposited as preprints (blue), COVID-19 related articles that were not deposited as preprints (red), and all non-COVID-19 articles published during the same period (light blue). Impact factors are displayed above the bars and are from the Journal Citation Reports [71].

**Table 6.**
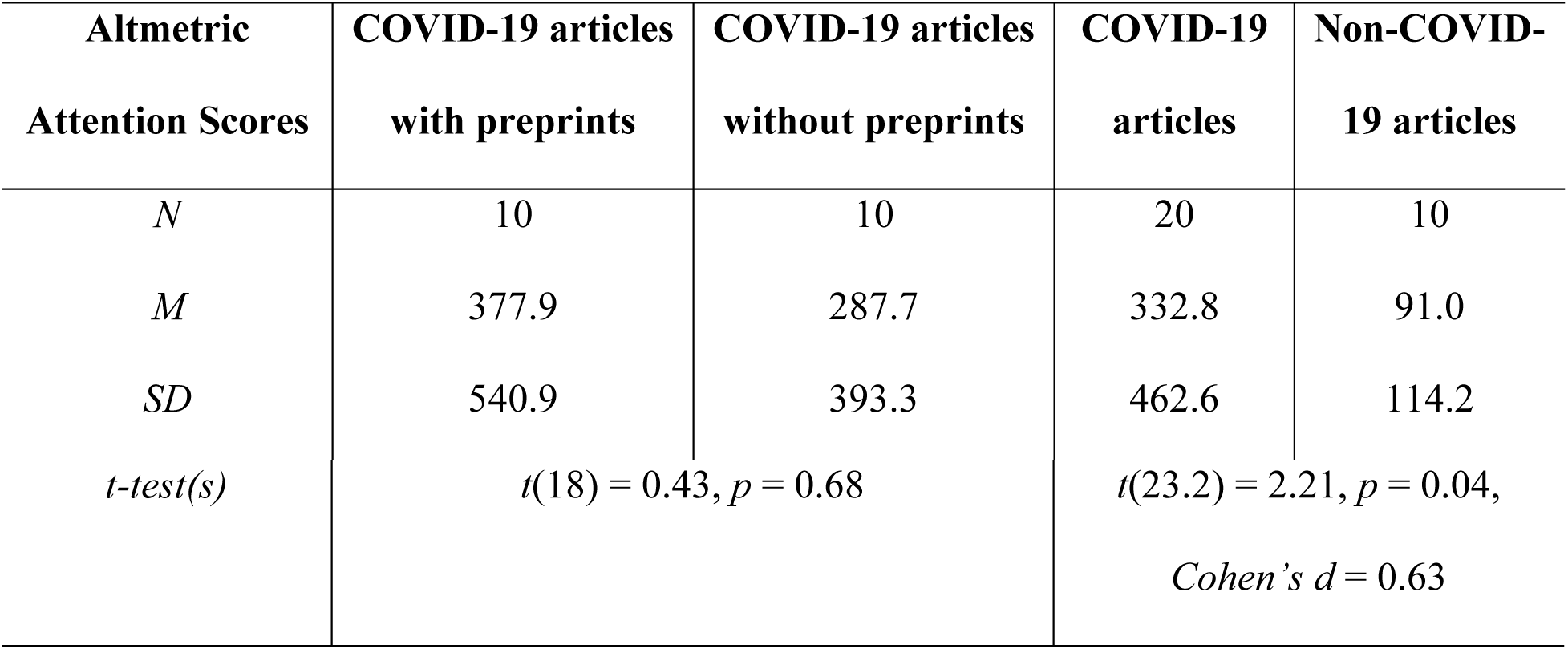
Descriptive statistics for Altmetric Attention Scores for COVID-19 related and unrelated publications, as well as Student’s t-test to evaluate discrepancies.

## Discussion

In this paper, we explored how publication practices in biomedical sciences reacted to an emergency, such as COVID-19 pandemic. Our first focus was analyzing the usage of two major biomedical preprint servers, bioRxiv and medRxiv. Following the deposition of the first preprint on a “novel coronavirus” in mid-January 2020 [56], preprint submissions to these two platforms increased rapidly. Submissions of new coronavirus related preprints reached 10 to 20 per day by February and increased to about 150 per day by May (Fig 1). In addition to this incredible flow of COVID-19 preprints, we observed about a 35% increase in deposition of preprints on all other topics. Towards the end of 2020, the amount of posted COVID-19 related preprints has been declining each month (Fig 1) but preprints unrelated to coronavirus continue to be submitted at a relatively constant rate (Fig 2). Preprints traditionally represented only a small fraction of scholarly literature; however, during the pandemic the fraction of combined medRxiv and bioRxiv COVID-19 preprints over all journal and review articles indexed in PubMed in the given month increased from 2% in February to 15% in May (Fig 3). A higher flux of biomedical preprints was also observed for Zika and Ebola outbreaks [60], which was suggested to result from an increase in research activities, and the statement of prestigious journal publishers and funders on the importance of preprints and data sharing in public health emergencies [73]. In contrast, COVID-19 pandemic was accompanied by a severe reduction in research activities due to social distancing policies and stay-at-home orders issued around the world. We hence raised the question regarding the origin of the amplified usage of preprint servers during the COVID-19 pandemic?

The answer to this question lies within the trends in the most active fields in each preprint server. Our analysis revealed that the majority of COVID-19 related preprints in bioRxiv were deposited in those fields that are most relevant to coronavirus research, such as *microbiology*, *bioinformatics*, and *immunology* (Fig 4A); while the leading categories in all other topics during the same period were *neuroscience, microbiology*, and *bioinformatics*. Note that prior to the pandemic, the three leading categories in bioRxiv were *neuroscience, bioinformatics*, and *genomics* [51]. Thus, while *neuroscience* researchers remain as the main users of the bioRxiv platform, we see an emergence of a new community of preprint authors working in the fields of *immunology* and *microbiology*. The latter can perhaps be further refined as *virology*, a category currently absent in bioRxiv (and medRxiv), but listed among journal scope categories, where the majority of *microbiology* preprints were later published. With regard to medRxiv server, the leading categories for COVID-19 related preprints are *infectious diseases*, *epidemiology*, and *public and global health* (Fig 4B); however, it is not possible to compare these figures with the pre-COVID-19 data due to the relatively recent launch of this preprint server. To explain, medRxiv was founded only six months prior to the pandemic and increased its volume by almost 16-fold during 2020, with 66% of this increase attributed to preprints on COVID-19. Overall, we noted that during the coronavirus pandemic preprint servers attracted a new pool of researchers from previously underrepresented fields. This agrees well with an earlier report of Coates *et al.* who found that 83% of COVID-19 preprint authors were posting a preprint for the first time [62]. It is worth noting that the same study also found that most corresponding authors were not switching to COVID-19 research from other fields, therefore, we propose that the most plausible factor contributing to the observed surge of preprints is the extensive use of preprint repositories by new COVID-19 researchers. This is remarkable considering preprints are rarely taken into consideration for tenure or promotion decisions in biomedical sciences. Further studies monitoring these new clients of preprint servers will be valuable in determining whether this is just a transient behavior triggered by COVID-19 or whether these practices will persist beyond pandemic duration.

Once we established that increased volume arises from new users, we further inquired whether, besides rapid dissemination of research results, they were attracted by any specific of the known benefits for preprint platforms. Among many, we focused on: (*i*) opportunity to share studies that are difficult to publish in traditional journals or those that would be rather works-in-progress than complete reports, and/or (*ii*) access to public comments that could help improving the manuscript prior to journal submission.

We first explored whether COVID-19 preprints were in-progress or complete works by analyzing how many of the COVID-19 preprints later appeared in refereed journals, and used that metric as an indicator for their suitability for publication. Our analysis showed that by Sept 30, 2020, only 18% of the bioRxiv and medRxiv preprints related to COVID-19 appeared as peer-reviewed journal publications. Anticipating a dependence of publication rate on the analysis timespan [51], we assessed the time it takes for a preprint to resurface as a journal article and found that our study on Sept 30, 2020, undervalued publication rate for preprints deposited any time after the end of July (mean *TΣ*= 63.4 days, Table 2). To derive a meaningful publication rate, we reanalyzed the same pool of COVID-19 related preprints (deposited to bioRxiv and medRxiv servers by Sept 30) on December 7, 2020, which is 68 days past Sept 30 (mean bioRxiv *TΣ*= 68.5 days, Table 2). As expected, we found higher publication rates for COVID-19 preprints, 33.8% and 28.8% for bioRxiv and medRxiv servers, respectively; although these values remained lower than those reported for pre-pandemic preprints (42% [51] or 70% [3]) and for previous health crises, such as Ebola and Zika (60% and 48%, respectively) [60]. We therefore concluded that COVID-19 preprints are mostly works-in-progress, in line with earlier suggestions [62]. In the future, it will be important to reevaluate publication rates for COVID-19 related preprints. It is likely that some COVID-19 preprints keep circulating through repeated cycles of journal submissions and rejections, and thus remain invisible to our analysis.

We then assessed whether coronavirus researchers used bioRxiv and medRxiv preprint servers to gather public feedback by examining the *pre-submission time* (*tα*) for COVID-19 preprints. Our analysis of publication delays yielded a median *tα*of 0 days for published COVID-19 preprints, implying that preprints were submitted to preprint servers and to journals simultaneously (Fig 9 and Fig 11). A more detailed analysis showed that only 28% of preprints were deposited into servers for over 10 days prior to journal submission. This trend of shortening the waiting period between preprint deposition to a server and its submission to a journal had been already reported in studies involving bioRxiv [69] and arXiv preprints [70]. Note that *pre-submission time* may include submissions to multiple journals, thus a low *tα*value also means that the majority of the so far published COVID-19 preprints were accepted to journals at their first submission. Thus, we see no change in authors’ behavior due to pandemic: COVID-19 manuscripts deposition as preprints occurred concurrently to their submission to journals; this implies that authors of COVID-19 preprints did not specifically pursue the pre-submission feedback when posting their research study on a preprint server.

In line with previous reports [52,61,67], our analysis of publication delays demonstrated another important change in scientific publishing in response to the COVID-19 pandemic, namely, expedited publication process for COVID-19 related articles. For instance, COVID-19 article versions of preprints are currently reviewed in about half the time (43 days) as compared with the 30-year average *peer-review time* of 100 days [65]. Further, COVID-19 articles associated with preprints are being reviewed 70% faster and prepared for the final version 90% faster than biomedical articles were prior to the pandemic (*t_R_* = 141 days and *tβ*= 147 days, in 2013 [66]). This acceleration is also evident when comparing articles associated with bioRxiv preprints: COVID-19 related preprints transformed into peer-reviewed journal publications 2.5 times faster during the study period, Jan 1 – Oct 23 2020, as compared to bioRxiv preprints before the pandemic (*T_Σ_* = 63.4 days in 2020 *vs* 155-166 days by 2018 [51, 64]; Table 3). This is unprecedented and was not observed for Zika or Ebola related preprints, where the median *elapsed time* for both outbreaks (150 days [60]) was reported to be no different than the normal publication timeline.

This really speaks to a success of modified journal editorial policies in expediting publication process for coronavirus researchers. These initiatives [33, 74] quickly led to a surge of COVID-19 journal articles that in July represented about 62% of all journal and review articles indexed in PubMed for that month. By the end of October 2020, 1,357 of COVID-19 preprints from medRxiv and bioRxiv servers appeared in 525 different academic journals, with multidisciplinary journals, such as *PLOS ONE*, *Science*, *Nature*, and *Nature Communications*, accruing the highest number of former preprints (see Leading Journals in SI). Among the specialized journals, *Clinical Infectious Diseases*, and *Journal of Medical Virology* published 6% of all medRxiv preprints. The COVID-19 portfolio of *BMC Medicine*, *Journal of Clinical Microbiology*, and *Nature Communications* was on one third composed of former preprints. However, the highest fraction of preprints in all articles published by the journal was observed for *Emerging Microbes & Infections*, where 69% of COVID-19 articles originated from preprints. Journal champions varied at various moments throughout the pandemic, which was found to be related to variations in publication times among the journals (Fig 12). For example, it took the fastest for bioRxiv or medRxiv preprints to appear in *Journal of Clinical Virology* or *Journal of Medical Virology* (*TΣ*= 26.2 and 38.3 days, respectively), but longest in *PLOS ONE* and *Nature Communications* (*TΣ*= 89.4 and 102.7 days, respectively). On average, the publication process for COVID-19 preprints posted to bioRxiv or medRxiv servers took about two months. This reflects an earlier observation that a publication peak for COVID-19 preprints in May transfers to the summit in July for journal article publications (Fig 2).

Complementary efforts of preprint servers and scholarly journals to disseminate knowledge promptly, while differentiating reliable and important findings from those that may be misleading attest to the upmost relevance of COVID-19 topic during 2020, as evident from Altmetric Attention Scores for COVID-19 research articles (Fig 14). For example, COVID-19 related articles published in *Nature* journal between Jan 1 and Nov 19, 2020 had a mean altmetric score of 1,077.8 as compared to 354.3 for articles on all other topics. The highest Altmetric Attention Score of 1,518.2 was observed for *Nature* journal articles associated with preprints. Despite the high visibility of COVID-19 articles originated from preprints, our analysis showed no significant altmetric advantage over COVID-19 articles that did not have a preprint version. This is in contrast with previous findings of clear altmetric differential between articles deposited as bioRxiv preprints and those that were not [18,64,71]. Considering the acute public interest in novel coronavirus research, we believe that any scholarly work, whether it was an article, associated or not associated with a preprint, received equivalent amount of social media attention during the pandemic. The strong correlation we observed between the altmetric scores and journals’ impact factors for COVID-19 articles tells us how popularized was the professional arena in biomedical sciences during the pandemic.

In summary, our analysis showed that early in pandemic, preprints were prevailing in disseminating findings on the topic of the public health emergency. Preprint authors deposited them into fields previously underrepresented on bioRxiv or medRxiv servers but those that were directly related to our understanding of the newly emerged coronavirus and ways to prevent the spread of the disease. This new category of authors mainly pursued rapid and transparent scientific communication but was not specifically interested in pre-submission feedback as observed from preprints submissions onto servers occurring concurrently with the journal submissions. We believe the majority of COVID-19 preprints were immediate in-progress findings, not suitable for the direct transfer to refereed journals; this conclusion was based on the low publication rate for COVID-19 preprints. The originally estimated publication rates were reaffirmed two months later, which is how long it took on average for a COVID-19 preprint to go through the peer-review and production stages. Both stages of the publication process were significantly expedited for COVID-19 publications; but they varied widely among different journals. The COVID-19 preprints that resurfaced as journal articles display exceptionally high Altmetric Attention scores echoing a high social media engagement in coronavirus research. The concerted efforts of journal and preprint publishers, institutions, funders, and individuals to bring us information and to facilitate its sorting and evaluation, deserve special applauds. It is our hope that as pandemic recedes, biomedical sciences will keep on building their relationships with preprint servers, while journal publishers will retain policies that were most helpful in expediting, revising, and sharing critical research output during the pandemic.

## Acknowledgements

We thank the University of Michigan Library for the financial support of this project and CADRE Fellowship from the Research Cohort for the Study of Coronaviruses Program for providing access to the CORD-19 dataset. We thank Dr. Craig Smith for statistics related consultations. We also thank Dr. Oscar Tutusaus for his assistance with manuscript editing.

## Supporting Information

### Supplementary files

1. Supporting information that contains:

a. Data Flow Chart – shows the usage of various data platforms in this study;
b. Published Collections – shows how various data were combined to account for all published preprints;
c. Leading Journals – shows the number and relative fraction of COVID-19 preprints published in selected journals and highlights leading journals in publishing COVID-19 bioRxiv and medRxiv preprints;
d. Publication Rates − shows publication rates for COVID-19 preprints.
2. CSV files with raw data for:

a. Altmetric Attention Scores – shows average altmetric scores and impact factors for selected journals;
b. Analysis of Journal Categories – shows how journal categories relate to preprint categories for published bioRxiv and medRxiv COVID-19 preprints;
c. Analysis of Preprint Categories − shows data on categories of bioRxiv and medRxiv preprints prior to and during the pandemic, both for COVID-19 related and unrelated works;
d. ANOVA – shows the results of ANOVA test to evaluate whether one parameters for one journal were significantly different from parameters for a group of journals;
e. Articles’ Publication Pates – shows the variation in publication dates retrieved from different data sources;
f. Publication Delays – shows various publication delays and their distribution for COVID-19 preprints published during 2020;
g. Scholarly Output – shows the monthly accumulation rates for bioRxiv and medRxiv preprints, as well as for journal and review articles related and unrelated to COVID-19, as indexed in PubMed;
3. Python code required to pull the data and visualize selected figures.

## Data availability

Source data for all figures have been provided in supporting files that were deposited in a Zenodo repository with DOI 10.5281/zenodo.4329576.

